# Inference of evolutionary transitions to self-fertilization using whole-genome sequences

**DOI:** 10.1101/2022.07.29.502030

**Authors:** Stefan Struett, Thibaut Sellinger, Sylvain Glémin, Aurélien Tellier, Stefan Laurent

## Abstract

The evolution from outcrossing to selfing is a transition that occurred recurrently throughout the eukaryote tree of life, in plants, animals, fungi and algae. Despite some short-term advantages, selfing is supposed to be an evolutionary dead-end reproductive strategy on the long-term and its tippy distribution on phylogenies suggests that most selfing species are of recent origin. However, dating such transitions is challenging while it is central for this hypothesis. We develop two methods making use of full genome polymorphism data to 1) test if a transition from outcrossing to selfing occurred, and 2) infer its age. The sequentially Markov coalescent based (*teSMC*) and the Approximate Bayesian Computation (*tsABC*) methods use a common framework based on a transition matrix summarizing the distribution of times to the most recent common ancestor along the genome, allowing to estimate changes in the ratio of population recombination and mutation rates in time. We first demonstrate that our methods can disentangle between past change in selfing rate from past changes in demographic history. Second, we assess the accuracy of our methods and show that transitions to selfing as old as approximatively 2.5*N*_e_ generations can be identified from polymorphism data. Third, our estimates are robust to the presence of linked negative selection on coding sequences. Finally, as a proof of principle, we apply both methods to three populations from *Arabidopsis thaliana*, recovering a transition to selfing which occurred approximately 600,000 years ago. Our methods pave the way to study recent transitions to predominant self-fertilization in selfing organisms.

**Significance statement:** Self-fertilization evolved recurrently from outcrossing in many groups of organisms. When, why, and at what pace such transitions occurred are central to understand the evolution of reproductive systems but dating them remains highly challenging. While experimental work can be conducted in ecological set-ups, it is difficult to reconstruct and empirically test the past ecological conditions which could have driven transitions from outcrossing to self-fertilizing reproduction. We suggest here to use full genome data of several individuals per population to estimate if and when a transition in reproductive strategy occurred. We develop two methods which can be applied to estimate the age of such transitions jointly with the species demographic history.

## Introduction

Hermaphroditism is common in many groups of eukaryotes, especially in plants, and allows uniparental reproduction through selfing. The rate of self-fertilization is known to vary widely between species and populations; from exclusive outcrossing, through mixed mating, to predominant self-fertilization (Whitehead, et al. 2018). In flowering plants, cross-fertilization is ensured by diverse molecular and morphological features, some of which being referred to as self-incompatibility (SI) mechanisms (Charlesworth 2010), which are defined as “the inability of a hermaphroditic fertile seed plant to produce zygotes after self-pollination” (de Nettancourt 1977). SI mechanisms enable the pistil of a plant to identify and repel self-pollen or pollen of a related genetic type and as a consequence, avoid inbreeding. SI systems are known to be encoded by a small number of genes and can therefore easily be lost. Indeed, genetic disruptions of SI systems through naturally occurring mutations are thought to be a major driver of plant reproductive diversity and have been linked to recent transitions to predominant self-fertilization in several species (Shimizu and Tsuchimatsu 2015; Mattila, et al. 2020). Transition from outcrossing to predominant self-fertilization is the most frequent reproductive transition in flowering plants (Barrett 2010) and is thought to have occurred many hundreds of times. It is one of the best studied reproductive transitions owing to its profound ecological, evolutionary, and developmental consequences affecting genetic and morphological diversity as well as patterns of dispersal. It also nicely illustrates how the balance between micro-evolutionary and macro-evolutionary processes can generate the observed distribution of mating systems among species (Igic, et al. 2008; Goldberg, et al. 2010).

On the short term, whether a new mutation responsible for a SI breakdown will invade the population or be lost depends on the balance between the two main advantages of selfing (reproductive assurance and gene transmission advantage) and inbreeding depression (the reduced fitness caused by an increased homozygosity under inbreeding (Charlesworth and Charlesworth 1987). On the long-term, selfing is predicted to increase extinction rate and to reduce diversification of selfing lineages as it has been observed in a few clades such as Solanaceae (Zenil-Ferguson, et al. 2019), Primulaceae (de Vos, et al. 2014); but see Landis, et al. (2018). A consequence is the supposed recent origin of selfing species due to an excess of transitions on terminal branches. This peculiar dynamic is also invoked to explain that the genomic effects of selfing are often detected on within-species polymorphism but very rarely on between-species divergence (Glémin, et al. 2019). The recent timing of these transitions is thus a central assumption but has not been systematically tested as it remains a challenging task.

Despite the increasing number of available methods to reconstruct the evolutionary history of populations using genomic data (Excoffier, et al. 2013; Schiffels and Durbin 2014; Boitard, et al. 2016; Speidel, et al. 2019), none of them is currently able to explicitly identify and estimate the age of transitions in reproductive strategies. In flowering plants, previous attempts to estimate the age of transitions to selfing were based on limited genetic variation at a single locus directly controlling the reproductive mode in plants: the S-locus (reviewed in Mattila, et al. (2020)). For example, naturally occurring loss-of-function alleles at the S-locus were shown to be responsible for the loss of self-incompatibility in *Arabidopsis thaliana* (Tsuchimatsu, et al. 2010); and the steady accumulation of non-synonymous alleles following loss of constraint at the S-locus was used to estimate that the age of the transition is at most 430,000 years old (Bechsgaard, et al. 2006). However, this approach is limited by the small number of genetic variants upon which the estimation is conducted and can only be used in species for which, as is the case in *A. thaliana*, the genetic determinism of the loss of self-incompatibility is known. However, a shift in reproductive system is expected to strongly impact genome-wide polymorphisms patterns, not only at the loci controlling it, thereby leaving a potentially characteristic molecular signature. We used this rationale to develop two inference tools allowing to use full genome polymorphism data of any species in order to 1) reveal the occurrence of past changes in reproduction mode, and 2) estimate their age.

A classic way to consider selfing assumes a theoretical population of *N* diploid individuals which produce offspring through selfing or outcrossing with probability *σ* and 1 – *σ*, respectively. Under a model of neutral evolution, the distribution of polymorphic sites in a sample of sequenced individuals, that is the frequency of alleles (single nucleotide polymorphism, SNPs), is determined by the underlying genealogy of this population. A genealogy has for properties its length measured as the time to the most recent common ancestor (*T*_MRCA_) and its topology, that is the order and number of branching processes. Along the genome, genealogies change due to the effect of recombination (the so-called ancestral recombination graph, ARG, (Hudson and Kaplan 1988; Wiuf and Hein 1999). Two population parameters determine the distribution and characteristics of genealogies observed in a sample of several genomes: the population mutation rate (*θ*) and the population recombination rate (*ρ*). In the presence of predominant selfing, the effective population size (*N*_σ_) of a population of *N* individuals is scaled by the selfing rate (*σ*) yielding *N*_σ_=*N*/(1 + *F*) (Nordborg and Donnelly 1997) and the recombination rate (*r*_σ_) is scaled as *r*_σ_=*r*(1 – *F*) (Nordborg 2000; Padhukasahasram, et al. 2008), where *r* is the recombination rate in the genome and *F* is the inbreeding coefficient. I*F* is defined as *F*=*σ*/(2 – *σ*) and takes values between 0 and 1 (0 for outcrossing, and 1 for fully selfing). As a consequence, the population recombination rate takes now for value in a selfing population: *ρ_σ_=*4*N_σ_ r_σ_=*4*Nr*(1 – *F*)/(1 + *F*) [1].

With *μ* being the mutation rate in the genome, the population mutation parameter accounts for the effect of selfing as follows: *θ_σ_* = 4*N_σ_μ = 4Nμ*/(1 + *F*) [2].

We note that the classic ratio of population recombination by population mutation rate *ρ/θ = r/μ* in outcrossing species, becomes now with selfing *ρ_σ_/θ_σ_ = r*(1 – *F*)/*μ* (Nordborg and Donnelly 1997; Möhle 1998; Nordborg 2000). Taken together, expressions (1) and (2) suggests that selfing does not amount to a simple change in effective population size (from *N* to *N_σ_*), and that the reduction of the effective recombination rate is more severe than the reduction in effective population size.

Following these insights, two key predictions can be derived. First, a characteristic and specific signal of selfing, in contrast to outcrossing, is expected to be present in SNP data due to the joint action of recombination along the genome (rate *ρ_σ_*) and of the genealogical (coalescence) process (rate *θ_σ_*). The first prediction underlies the previous development of a sequentially Markov coalescent method (eSMC) to estimate a fixed selfing rate using estimations of the ratio *ρ_σ_/θ_σ_* from pairs of genomes (Sellinger et al. 2020). Second, evolutionary changes in reproductive systems (transition from selfing to outcrossing or *vice and versa*) are reflected in variations of the ratio *ρ_σ_/θ_σ_* in time and are identifiable and distinguishable from changes in population sizes alone. The latter suggests that a characteristic signature of the change in selfing rate in time should be observed in polymorphism data, if genetic variation can be summarized in a way that is informative about the joint effect of genetic drift and recombination. Indeed, it is desirable to 1) estimate changes in population size which occurs independently of a transition, for example when species colonize new habitats as facilitated by selfing, and 2) disentangle the possible confounding effect of population size changes on the estimation and detection of a transition. To our knowledge no statistical inference method exists that takes advantage of these theoretical predictions to jointly estimate temporal changes in selfing rates and in population sizes. We first provide results on the consequences of a transition to selfing on genomic variation that confirm our second prediction. Then, we analyse the statistical accuracy of the two newly developed methods to identify and estimate the age of transitions to selfing using a small number of sequenced genomes. Third, as a proof of principle, we apply these methods to estimate the age of the transition to selfing in *Arabidopsis thaliana*, in which it has been documented and for which full genome polymorphism data exist. These new methods are welcome toolkits for dating and understanding the evolution of the selfing syndrome and shifts in breeding systems and reproduction modes.

## Results

### The consequences of a transition to selfing on patterns of genomic variation

We consider a theoretical population of *N* diploid individuals (equivalent to 2*N* chromosomes), which produce offspring through selfing or outcrossing with probability *σ* and 1 – *σ*, respectively (following the notations in Nordborg (2000)). At some time (*t*_σ_) in the past, the previously outcrossing population (with *σ* = 0) undergoes a transition to selfing and remains selfing until present (with a selfing rate *σ* > 0). Independently of the change in selfing rate, the population size can change once from *N*_ANC_ (ancestral) to *N*_PRES_ (present) at time *t*_N_ (measured from the present). We implement the selfing model both in the forward Wright-Fisher framework, in which selfing can be simulated explicitly, and using the coalescent-with-selfing, in which selfing is modeled through a rescaling of the effective population size and of the recombination rate at *t*_σ_ (Nordborg and Donnelly 1997; Möhle 1998; Nordborg 2000)(see methods).

We here verify the second prediction using simulated distributions of *T*_MRCA_-segments (hereafter TL-distribution). These segments are defined as successive and adjacent sets of nucleotidic positions sharing the same time to the most recent common ancestor (*T*_MRCA_) and are separated by the breakpoints of ancestral recombination events (McVean and Cardin 2005). Segments are summarized by their lengths and *T*_MRCA_ (Figure 1A, B). When the selfing rate is constant the TL-distribution has a negative constant covariance, because, on average, older segments are exposed, and shortened by a larger number of recombination events compared to younger segments (Figure 1A). In the case of a transition from outcrossing to predominant selfing, the rate at which the segments shorten with age is dramatically increased in the outcrossing phase compared to the selfing phase (Figure 1B); leading to a characteristic change in the covariance of the joint distribution at *t*_σ_. This behavior of the model is invariably observed, whether we simulate selfing by re-scaling *ρ* and *θ* or explicitly using forward Wright-Fisher simulations (Figure S1). Importantly, this specific genomic signature of a transition to selfing is not observed when the selfing rate is constant and only the population size changes from *N*_σ_ to *N*, where *N*_σ_ would be the effective population size of a population with selfing rate *σ* (Figure 1A). Thus, in the coalescent framework, when selfing rates are not constant, the probability of a recombination event is not increasing linearly with time, but also depends on the selfing values at each time point (Figure S2).

**Figure 1.**
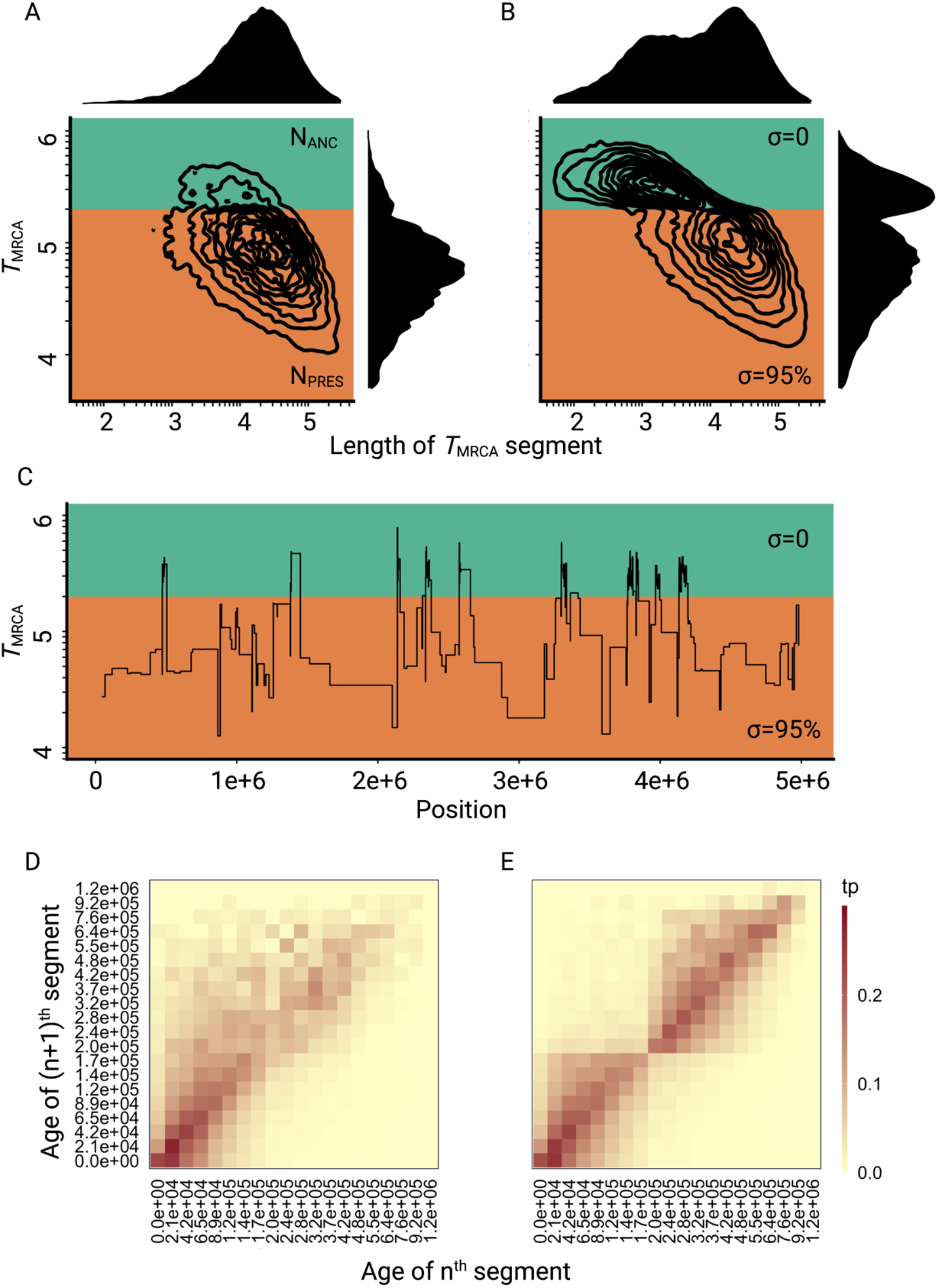
Consequences of a transition to selfing on the genealogies of simulated chromosomes. **A:** Joint and marginal distributions of ages (*T*_MRCA_ in generations on a log10 scale) and lengths of *T*_MRCA_-segments (in bp on a log10 scale) in a selfing population (*σ*=0.95) with a stepwise change from large (green, *N*_ANC_ = 50,000) to low (orange, *N*_PRES_=26,250) population size. The population sizes were chosen to correspond to the rescaling of the effective population size by the selfing rates used in panel B. **B:** Joint and marginal distributions of ages (*T*_MRCA_) and lengths of *T*_MRCA_-segments (in bp) in a population with a constant population size and a shift from outcrossing (green, *σ*=0) to predominant selfing (orange, *σ*=0.95). **C:** Spatial distribution along the genome of a subset of the *T*_MRCA_-segments simulated in panel B (*t*_α_=200,000 generations). **D:** The transition matrix of ages (*T*_MRCA_) between adjacent segments along the genome for the data simulated in panel A. This matrix summarizes the probabilities that the *n*^th^ *T*_MRCA_-segment with a given age *X* is followed by the (*n*+1)^th^ segment of age *Y*. The heat colors indicate the transition probabilities (tp). **E:** The transition matrix of ages (*T*_MRCA_) between adjacent segments along the genome for the data simulated in panel B. The recombination rate for the simulations was set to 3.6×10^-9^ (see methods).

Although the same increase in *ρ* could in principle be accounted for by a very large ancestral population size (i.e. *N*_ANC_=*N*_PRES_/(1 – *F*)), such a model would also have a much larger ancestral *θ* compared to the selfing model, and thus cannot produce the same TL-distribution as a transition to selfing (and thus is not a confounding scenario for a transition to predominant self-fertilization). Simulated TL-distributions under a range of values for *t*_σ_ illustrate the dependency of the change in the covariance between the age and length of *T*_MRCA_-segments on the age of the transition (Figure S3). Thus, this suggest that genome-wide polymorphism data contains information about shifts to selfing, when the age of the transition falls well within the distribution of *T*_MRCA_.

We also made the important observation that all the segments that coalesce in the outcrossing phase, trace back their ancestry to a subset of segments that do not coalesce more recently than *t*_σ_, a constraint that is responsible for the observed spatial clustering of segments with *T*_MRCA_ older than *t*_σ_ along simulated chromosomes (Figure 1C). This effect can also be captured by inspecting the transition frequencies of *T*_MRCA_ for a large number of successive and adjacent fragments (Figure 1D, E). In the case of a transition to selfing, the *T*_MRCA_ transition matrix distinctly shows that segments with *T*_MRCA_ older than *t*_σ_ are more likely to be followed by segments that are also older than this time. Although a similar dependency between successive *T*_MRCA_ also exists when selfing is constant, the magnitude of the effect is more pronounced in the case of a shift to selfing. A more formal description of this effect in the context of the sequentially Markov coalescent is provided in Appendix A. The specific genomic signature of a transition to selfing and its equivalence to two simultaneous temporal changes in effective recombination rate and population size motivated us to design two new statistical methods to estimate the age of transitions to selfing.

### Statistical methods to estimate the age of a transition to selfing: teSMC

The distributions of *T*_MRCA_ along a pair of chromosomes sampled from a sexually reproducing population can be modeled and approximated assuming the sequentially Markov coalescent (SMC) (McVean and Cardin 2005). SMC-based methods infer model parameters, such as demographic histories, from fully-sequenced genomes and have been implemented in several statistical software used to estimate changes in effective population size through time under the assumption of a Wright-Fisher model (Li and Durbin 2011; Schiffels and Durbin 2014; Terhorst, et al. 2017). Recently, a new SMC method, *eSMC*, has been developed to include the effect of seed-banks and self-fertilization in order to estimate these parameters (i.e. dormancy and selfing rates) jointly with past population sizes (Sellinger, et al. 2020). However, *eSMC* assumes the rate of selfing and dormancy to be constant through time. Therefore, we here extend *eSMC* into *teSMC,* allowing the estimation of varying selfing or recombination rates through time, jointly with varying population size. To do so, *teSMC* no longer assumes the ages of recombination events to follow a uniform distribution along the branch representing the recombining lineage (Li and Durbin 2011; Schiffels and Durbin 2014); but rather let them be a function of the selfing and recombination rate at each hidden state. This allows *teSMC* to jointly infer piecewise functions of selfing or recombination rates and population sizes through time (methods in Appendix A). In order to account for prior knowledge, two modes are implemented for parameter inference: 1) the free mode, in which each hidden state has its own independent selfing/recombination rate, and 2) the single-transition mode in which *teSMC* estimates only three parameters (the current and ancestral rates, and the transition time between both rates), a constraint greatly reducing the number of inferred parameters and well suited for the analysis of recent and sudden shifts from out-crossing to predominant self-fertilization.

First, to demonstrate the theoretical accuracy of our model and inference method, we analyze its performance when sequences of *T*_MRCA_ are given as input instead of sequence data. This is termed the best-case convergence of *teSMC* (Sellinger, et al. 2021). We simulate data from a population undergoing a strong bottleneck and simultaneously a transition to selfing or change in recombination rate. In both cases the population size and the past selfing or recombination values are recovered with high accuracy (Figure S4). Second, to understand the convergence properties of *teSMC*, we analyze its performance under a simple scenario assuming a constant population size and a constant selfing value of 0.9 given different amount of data. We compare the *eSMC* method, which estimate a constant rate of selfing in time with *teSMC,* which estimates varying selfing through time (Figure S5). When selfing is known to be constant (*eSMC*), the value of this parameter is recovered with high accuracy and low variance even with the lowest amount of given data (Figure S5A). However, when it is not known whether selfing changes through time (*teSMC*), a greater amount of data is required to reduce the variance in the estimation (Figure S5B).

We now evaluate the statistical accuracy of *teSMC* on polymorphism data from genome pieces of 5Mb, simulated under a model with constant population size (*N* = 40,000) with mutation (*μ*) and recombination (*r*) rates of 1×10^-8^, and with an instantaneous change from outcrossing (*σ*_ANC_ = 0.1) to predominant selfing (*σ*_PRES_ = 0.99) at time *t_σ_* (see methods). Both the single-transition mode and the free mode estimation procedures performed well over the complete range of *t*_σ_ values (Figure 2A). Population sizes estimated under the assumption of a constant selfing rate are consistently larger than the true value in the outcrossing phase and display large fluctuations in the selfing phase, which could be mistaken for past population size bottlenecks (Figure S6). However, when *teSMC* is allowed to account for the change in selfing rate, population sizes estimates (*N*) remain close to the true values. We note that the increased variance in *N* in the selfing phase are likely caused by a smaller number of available *T*_MRCA_ segments.

**Figure 2.**
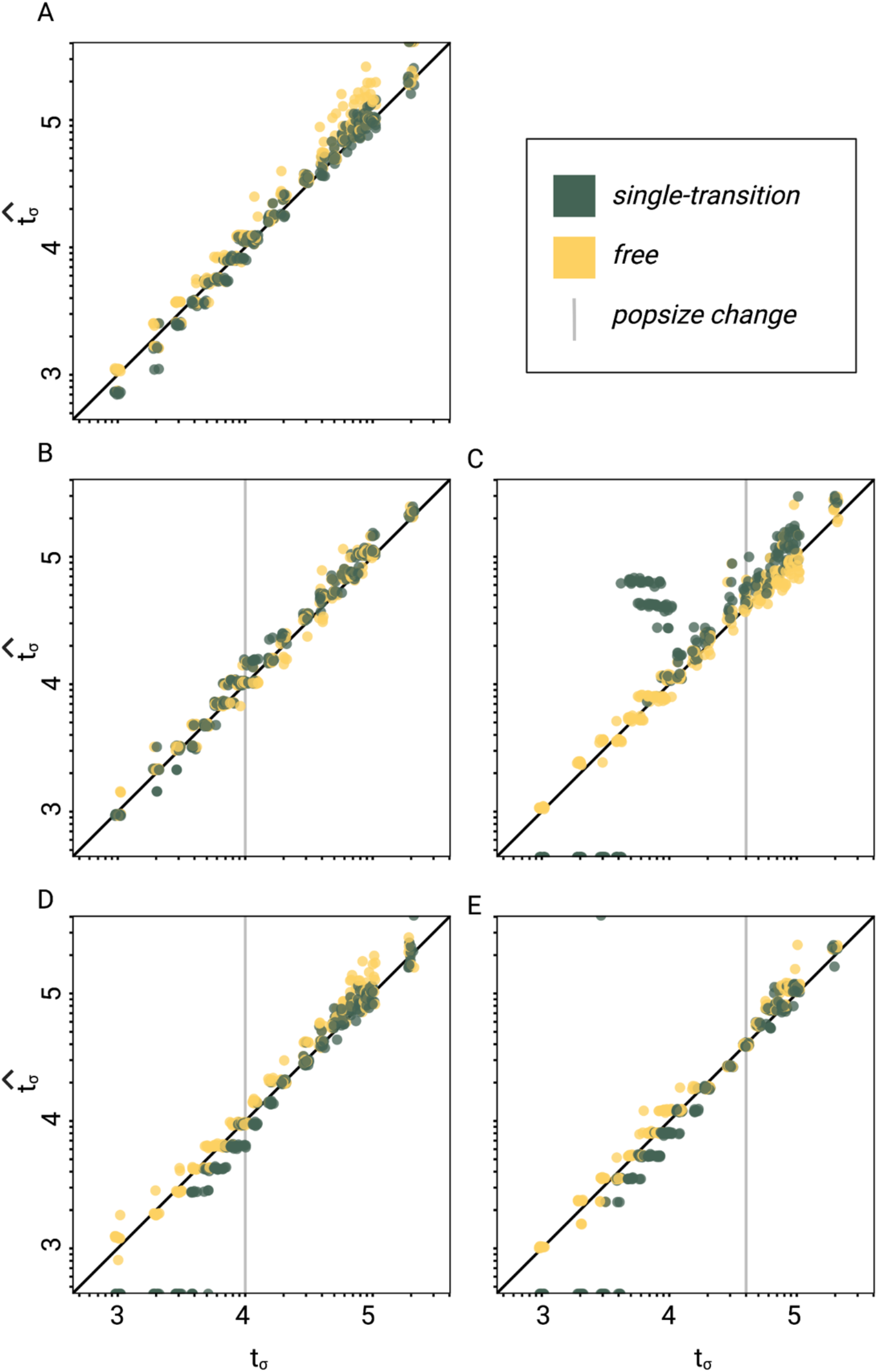
Performance of *teSMC* on simulated polymorphism data. Inference of times of transition from outcrossing (*σ* = 0.1) to predominant selfing (*σ* = 0.99) using neutral simulations. Inference was done using the free mode (yellow) and the one-transition mode (green) of *teSMC* and 10 replicates per time point. **A.** Under constant population size. **B-E:** simulations were done with an additional change in population size, the vertical grey line indicates the age of the change in population size. **B-C)** From *N*_ANC_ = 200,000 to *N*_PRES_ = 40,000 (population crash) at 10,000 generations (B) or 40,000 generations (C) in the past. **D-E)** From *N*_ANC_ = 40,000 to *N*_PRES_ = 200,000 (population expansion) at 10,000 generations (D) or 40,000 generations (E) in the past.

Finally, we evaluate the ability of *teSMC* to jointly estimate the age of a transition to predominant selfing and the time of a stepwise change in population size. To do this we use simulated data produced as above, except with the addition of a single stepwise population size reduction (Figure 2B,C) or expansion (Figure 2D,E). In both cases our results indicate that *teSMC*, especially the free mode inference method, is able to precisely estimate the age of the shift to selfing, regardless of the relative timing of the population size change and the transition to selfing. Also, in most cases, the population sizes inferred by *teSMC* were close to the true simulated values (Figures S7). However, when the transition is recent and the present population size is low, this can affect the precision of the population size estimates (Figure S7O). We note that *teSMC* fails to recover the population size, suggesting a lack of data (coalescent events) (Sellinger, et al. 2021). These results demonstrate that transitions to predominant self-fertilization and more generally large changes in recombination rate through time can be captured by *teSMC* and the estimations can be disentangled from changes in population sizes.

### Statistical methods to estimate the age of a transition to selfing: tsABC

In addition to *teSMC*, we develop *tsABC*, an approximate Bayesian computation (ABC) method to jointly estimate a change in population size and in selfing rate. ABC is a computational approach to estimate posterior probabilities for models and parameters that is well suited for demographic modelling in population genetics, where models often have many parameters and no analytically derived likelihood function (Beaumont, et al. 2002; Csillery, et al. 2010). Two advantages of the ABC method are that it allows to compare competing demographic hypotheses on basis of Bayes factors and it does not require bootstrapping the data to generate measures of uncertainty for the inferred parameters. A critical aspect of ABC is that it requires a careful summarization of the genomic data into a set of summary statistics that carry information about the parameters of interest (Beaumont, et al. 2002). In the case of a transition to selfing, we require that such summary statistics need to be informative about coalescence and recombination rates in order to make changes in selfing rates and population size distinguishable by the ABC model choice (Figure 3). Unfortunately, while the lengths of *T*_MRCA_-segments are straightforward to calculate on simulated genealogies (Figures 1A,B), it is more difficult to estimate them based on genomic diversity data alone. We calculate here the number of differences for pairs of sampled chromosomes using non-overlapping genomic windows of 10kb and construct a transition matrix for pairwise diversity similar to the one in Figures 1D,E; a summarization we refer to as TM_win_ (Figure S8, see methods). The rationale behind this method is that, if the window size is smaller than some of the clusters of *T*_MRCA_-segments in the outcrossing phase, then, TM_win_ will capture both the expected excess of diversity of those segments as well as their clustering (Figure 1C). For comparison we also considered classic summarization based on the site frequency spectrum (SFS) and a discretized distribution of the decay in linkage disequilibrium (LD-decay), as these carry information about temporal changes in population size and selfing rates (Tang, et al. 2007; Boitard, et al. 2016). We therefore evaluated the efficiency of three sets of summary statistics: SFS/LD, TM_win_, and SFS/LD/TM_win_ (see methods).

**Figure 3.**
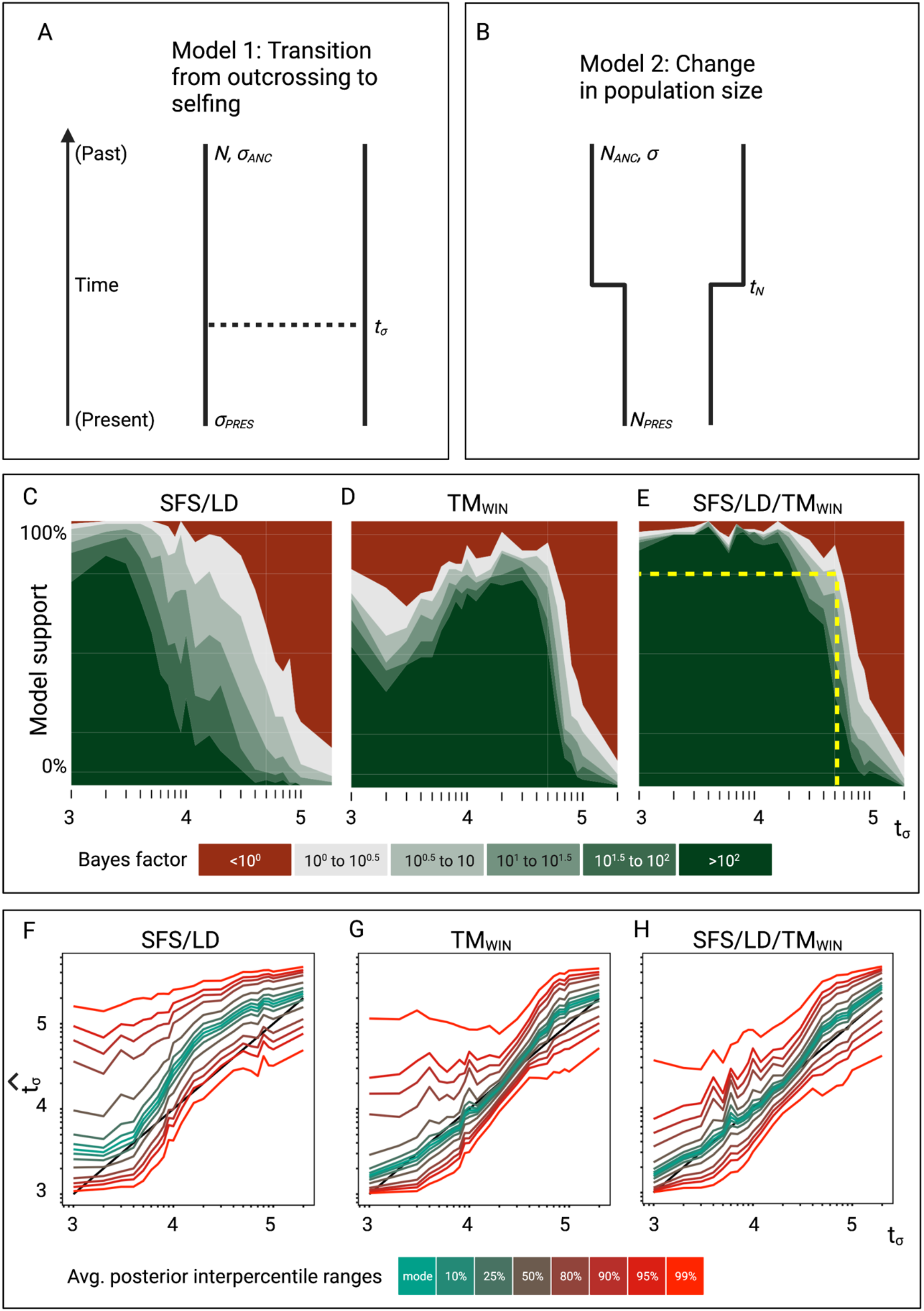
ABC model choice performance analysis. **A**: Demographic model 1 in the model choice analysis: one population with a single transition from predominant selfing to predominant outcrossing **B:** Demographic model 2 in the model choice analysis: one population with constant selfing and a single change in population size. The parameters of interest are the population sizes (*N*_PRES_, *N*_ANC_), the selfing rates (*σ*ANC, *σ*PRES), and the time of change in selfing rate and size (*tσ*, *t*N). **C, D, E**: Performance of the ABC model choice method using three different summarizations of data. **C**: Combining site frequency spectrum (SFS) and linkage disequilibrium (LD). **D:** Window-based transition matrix (TMwin). **E:** The combination out of SFS, LD and TMWIN. The x-axis represents the investigated range of *tσ* values; the y-axis indicates how often (out of 100 trials) *tsABC* correctly identified the transition-to-selfing model. The yellow dashed lines indicate that, for a Bayes factor of approximately 3 (BF=√10), *tsABC* identifies the right model 80% of the time, for transitions as old as 51,000 generations (corresponding to 2.5*N*e generations in coalescent time units, where *N_σ_* is the effective population size of the selfing population). **F, G, H**: Parameter estimation accuracy for the age of a transition to selfing (100 simulated datasets) under a model with constant population size (*N*= 40,000) and a change in selfing rate from *σ*ANC = 0.1 to *σ*PRES = 0.99. Coloured lines represent average quantiles for 100 posterior distributions.

To test whether our approach can distinguish a transition to selfing from a reduction in population size, we conduct an ABC model choice analysis using two competing models (Figure 3A,B): model 1 where the selfing rate changes from *σ_ANC_* to *σ_PRES_* at *t_σ_* and with *N* constant, and model 2 with population size changes from *N*_ANC_ to *N*_PRES_ and *σ* constant. We simulate datasets under model 1 and evaluated the ability of *tsABC* to identify the correct model for transitions of varying ages, using different sets of summary statistics to summarize the genetic data (see Appendix B). Our results show that our method recovers the correct model for transitions as old as 2.5*N_σ_* generations (Figure 3C-E), with *N_σ_* = *N*/(1 + *F*) being the effective population size for a selfing population. Interestingly, summarizing the genetic data using the SFS and LD-decay yields better performance for recent shifts to selfing (Figure 3C,D), while using TM_win_ performs better for shifts occurring between 0.5*N_σ_* and 3*N_σ_*); such that combining both set of summary statistics yields the best performance. This analysis confirms that shifts to selfing as old as approximatively 2.5*N_σ_* can be detected and can be disentangled from changes in population size using the ABC model choice procedure if appropriate summary statistics are used.

Next, we evaluate the accuracy of our method for estimating the age of a transition to selfing (*t_σ_*). We simulate 100 datasets under model 1 (Figure 3A) with values of *t_σ_* ranging from 1,000 to 200,000 generations, and used *tsABC* to re-estimate posterior distributions for *t_σ_* and the other parameters of the model (Figure 3F-H, Figure S9). Estimations are obtained using the same three summarization strategies used for the model choice (SFS/LD, TM_win_, SFS/LD+TM_win_). The age of a shift to selfing could be well estimated using the TM_win_ approach, while the SFS+LD approach over-estimated *t_σ_* almost over the complete range of values (Figure 3F,G). Combining SFS, LD, and TM_win_ does not further improve the accuracy of the estimations (Figure 3H). We note that the parameters *N* and *σ*_PRES_ (*i.e.* the population size and the current selfing rate) are both better estimated with TM_win_ than with SFS/LD, except for transitions younger than 10^4^ generations ago where *σ*_PRES_ is slightly better estimated with SFS/LD, Figure S9D). However, no set of summary statistics could estimate the ancestral selfing rate (Figures S9G-I).

### Robustness of inference to model violations

We here provide two analyses to demonstrate the robustness of our inference method to two violations of the model assumptions: first recombination and mutation rates may vary along the genome, and second, background selection in combination with selfing may affect the inference of transition to selfing. First, to assess the potential limits of our approach, we analyze the performance of *teSMC* when mutation and recombination rates are potentially non-constant along the genome (Figure S10-S11). When mutation and recombination rates are constant along the genome, *teSMC* recovers a constant population size and accurate selfing rates (Figures S10A, S11A). When recombination rate varies by a 2-fold factor along the genome, the estimation of population size is accurate (Figure S10B) but the variance of selfing rates inference increases (Figure S11B). Variation of the mutation rate by a 2-fold factor along the genome bias inference through time, leading to erroneously high inferred selfing rates and population size (Figures S10-S11 C,D). Yet, these results are reassuring as they demonstrate that no spurious transition from outcrossing to selfing in the past is inferred under variation of the recombination rate or mutation rate along the genome.

Background selection (BGS) refers to the effect of deleterious alleles on linked neutral diversity (Charlesworth, et al. 1993; Irwin, et al. 2016). Recently, several studies highlighted that neglecting the effect of BGS in demographic analyses can lead to statistical biases and potential miss-identification of population size changes (Ewing and Jensen 2016; Johri, et al. 2021). Because transitions to selfing result in strong reduction of the recombination rate (up to two orders of magnitude for transitions to predominant selfing), a corresponding increase of linkage between deleterious and neutral alleles is expected. Selfing indeed drastically magnifies the effect of BGS(Roze 2016). Because both the *teSMC* and the *tsABC* methods ignore the effect of selection, we evaluate their performance when applied to data simulated under a model with both a transition to selfing and background selection. We use *slim3* (Haller and Messer 2019) to simulate genomic data with the same distribution of exonic sequences as in five pre-defined regions of 1Mb from the genome of *A. thaliana* (exact coordinates in methods) and model negative selection on exonic sequences according to the distribution of fitness effects (DFE) published by Hamala and Tiffin (2020). We found that when exonic sequences are excluded from the analysis (masked), the accuracy to estimate the transition to selfing by *teSMC* improves slightly compared to the unmasked case (Figure 4A). The *tsABC* estimations remain accurate even without masking exonic regions (Figure 4B, Figure S12). These results suggest that our approach is generally robust to the effect of negative selection on linked neutral sites, even in compact genomes such as the one of *A. thaliana* (Figure 4).

**Figure 4.**
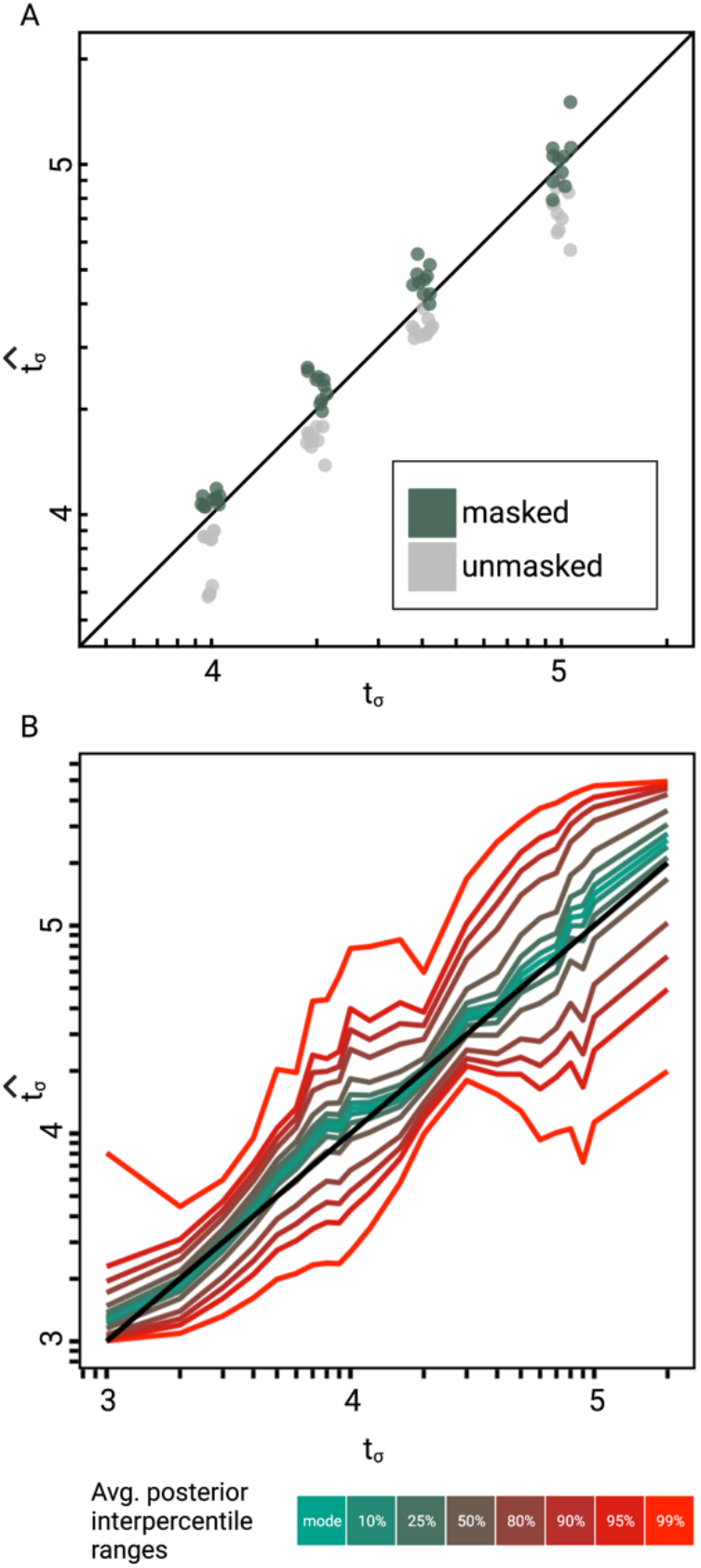
Accuracy of *teSMC* and *tsABC* in the presence of background selection (BGS). Inference of times of transition from outcrossing (*σ*=0.1) to predominant-selfing (*σ*=0.99) using **A)** *teSMC* and **B)** *tsABC*. Simulations were done under constant population size and negative selection acting on exonic sequences. The spatial distribution of exonic sequences was fixed and taken from the annotation of *A. thaliana*. Negative selection was modelled using the distribution of fitness effects from Hamala and Tiffin (2020) **A.** Comparison between simulated values of *t*σ and estimates obtained with *teSMC* using the one-transition mode. Estimations were conducted with and without masking exonic sequences subject to negative selection. **B**: Same analyses as in panel A but conducted with *tsABC*. Except for selection, simulations were done as in Figure 3H. Coloured lines represent the average quantiles for 100 posterior distributions obtained with *tsABC*.

### Application to Arabidopsis thaliana

Deactivation of the self-incompatibility mechanism through mutation knocking-out the gene *SRK* is known to be responsible for a transition to predominant self-fertilization in the model species *A. thaliana* (Tsuchimatsu, et al. 2010). This shift to selfing is the focus of several studies and estimates of its age vary widely depending on the type of data and statistical approach. Bechsgaard, et al. (2006) estimated the upper boundary for the age of the transition to be 413,000 years old based on phylogenetic analyses of the *S*-locus in *A. thaliana* and *A. lyrata*. Tang, et al. (2007) analysed the genome-wide decay in linkage disequilibrium but could not detect the expected signature of a transition to selfing, and therefore concluded that the shift must have been older than the oldest coalescent events in their sample (older than one million years approximately). Here, estimations obtained with *teSMC* and *tsABC* range from 592,321 to 756,976 depending on which method and population samples are used (Figure 5A, Table 1). Our estimates are older than the one proposed by Bechsgaard, et al. (2006) but younger than the age proposed by Tang, et al. (2007). Note that we are also able to jointly estimate the demography of each *A. thaliana* population and the transition to predominant self-fertilisation (Figure 5B). Furthermore, the estimates of transition to selfing are robust to the geographical origin of the population samples (Iberian non-relicts, Iberian relicts or central European, Figure 5).

**Figure 5.**
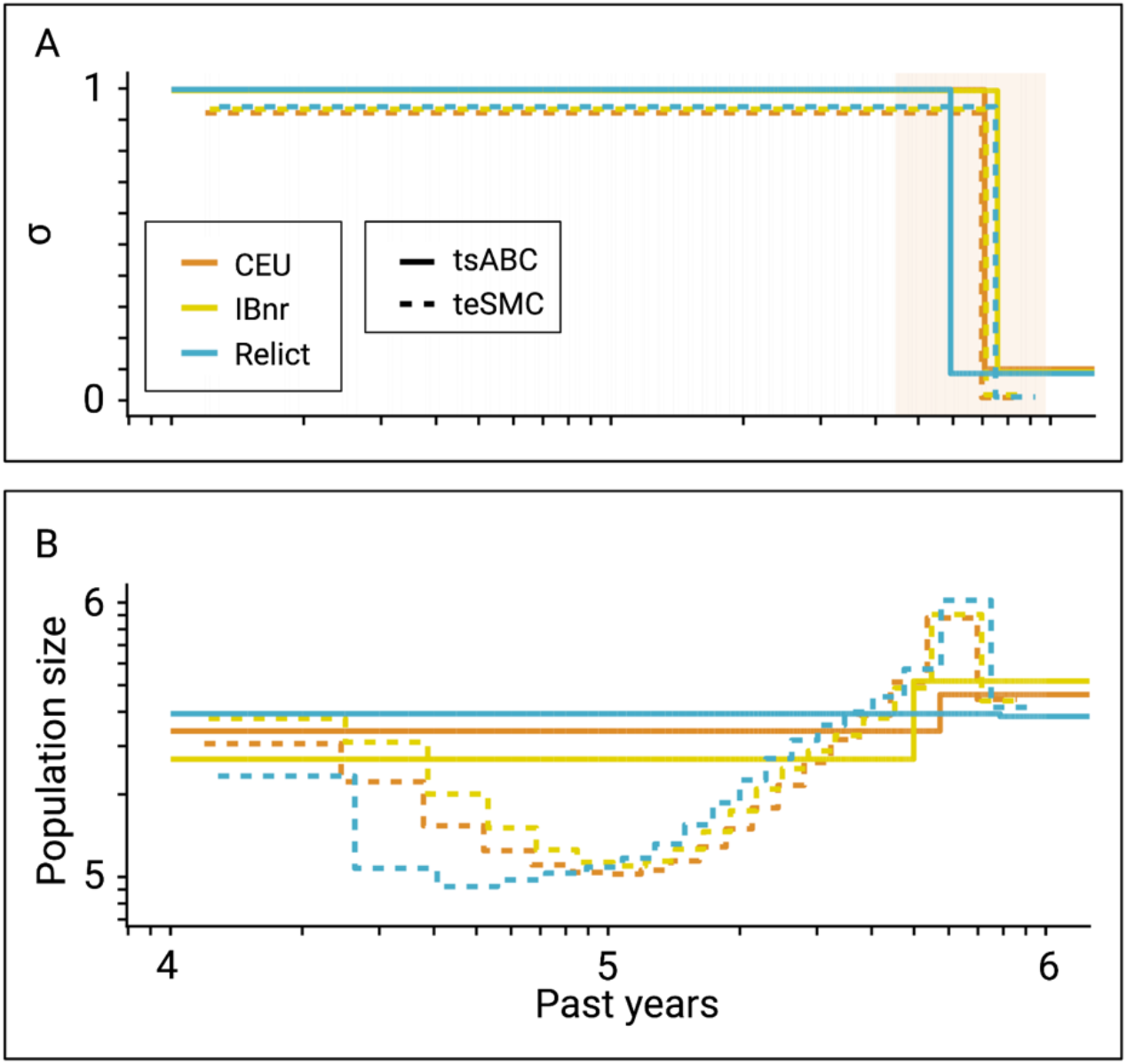
Inference of the time of transition from outcrossing to selfing in *Arabidopsis thaliana*. **A**: Inferred transitions from outcrossing to selfing for three independent genetic clusters of *Arabidopsis thaliana* from the 1001 genomes project (CEU, IBnr, Relict) using *tsABC* and *teSMC*. The 95%-CI (CEU) of the posterior distribution of *tσ* (*tsABC*) is indicated in light-orange. **B:** Co-estimated population sizes over time with a single population change (*tsABC*) or piecewise constant (*teSMC*).

**Table 1.** Estimated times of transitions from predominant outcrossing to predominant selfing in *Arabidopsis thaliana.* Estimations were conducted for three different ancestry groups: central Europe (CEU), Iberian non-relicts (IBnr), and Relicts using both *teSMC* and *tsABC*. For both methods, polymorphism data was measured on five genomic regions of 1Mb, located on the five chromosomes of *A. thaliana*. For *teSMC* exonic regions were excluded from the analysis. Only accessions with cluster membership > 95% were included.

| *Method* | *Population* | *Sample size* | *Mode* | *95% credibility interval* |  |
| --- | --- | --- | --- | --- | --- |
| *teSMC* | CEU | 20 | 697,490 | NA | NA |
| *teSMC* | IBnr | 20 | 713,421 | NA | NA |
| *teSMC* | Relicts | 17 | 749,668 | NA | NA |
| *tsABC* | CEU | 99 | 707,995 | 443,486 | 973,841 |
| *tsABC* | IBnr | 66 | 756,976 | 397,049 | 988,708 |
| *tsABC* | Relicts | 17 | 592,321 | 386,406 | 934,499 |

## Discussion

While the biological importance of shifts in mating systems or reproductive modes is long recognized as a key evolutionary and ecological process, no method based on genome-wide polymorphism data was available to estimate the age of a change in reproductive mode, while accounting for demographic history. In this study we show that by accounting for both the frequency distribution of SNPs and the distribution of historical recombination events, it is possible to recover shifts in reproductive modes based on a small number of fully sequenced genomes. The key idea underlying our approach is to leverage the properties of genomic segments delimited by recombination events (*T*_MRCA_-segments). Our simulations show that the molecular signature of a transition to selfing is two-fold. The first effect is a change in the relation between the age and the length of genomic segments that occurs at the time of the transition (Figure 1, S1-S3), and the second is a characteristic spatial distribution of the segments along the chromosome, where segments older than the transition tend to occur as clusters (Figure 1C-E). Several SMC-based methods already take advantage of both the distribution of age and lengths of *T*_MRCA_-segments to estimate past demographic history (Li and Durbin 2011; Barroso, et al. 2019; Steinrucken, et al. 2019), spatial structure (Wang, et al. 2020), heterogeneous recombination rates along the genome (Barroso, et al. 2019), or life-history traits such as dormancy (Sellinger, et al. 2020). SMC-based as well as other inference methods (Kerdoncuff, et al. 2020) rely on estimating changes in the ratio of population recombination rate by population mutation rate along the genome (the ratio *ρ/θ).* Yet, none of them attempted to leverage the information on the distribution of *T*_MRCA_-segments to also estimate temporal changes of the ratio *ρ/θ*, which in our case amounts to estimate variation of the selfing rate. Interestingly, as most inference methods are developed with a main application to human data in mind, the estimation of a changing *ρ/θ* ratio has not yet been a relevant and pressing question. Deng, et al. (2021) described how temporal recombination rates could be estimated from reconstructed ARGs, but only evaluated the performance of this approach on simulations with a constant rate.

In this study we develop two approaches to estimate change in selfing in time: one building upon the Markovian assumption of coalescence events along the genome (*teSMC*) and another one by considering the dependencies between the genetic diversity observed in successive genomic windows of fixed size (*tsABC*). The advantage of the *teSMC* approach is that it is most effective in identifying the boundaries and the ages of *T*_MRCA_-segments and therefore allows capturing the characteristic effect of a shift to selfing (as shown in Figures 1A, B). *teSMC* is also efficient in the way it optimizes the likelihood such that the method can be executed on a single desktop computer when applied to empirical data. The drawback of the *teSMC* method is that extending it to more complicated demographic scenario is mathematically and computationally difficult (*e.g.*, to selection or admixture). Conversely, the *tsABC* approach can easily be extended to more complicated demographic and selective scenarios but suffers from a lower statistical performance caused by the approximation made in the summarization of the data; and by the requirement to conduct many simulations that can only be realistically obtained on a high-performance computing cluster. Our results suggest our approaches to be robust to the presence of background selection, and to a lesser extend to variation of recombination and mutation rate along the genome. We nevertheless show that a spurious change in selfing rate is not generated by variation in recombination rate along the genome (Figure S11). We finally highlight the novelty in the design of the *tsABC* approach, which introduces a new summary statistic, the transition matrix for heterozygosity levels, which appears to contains similar information as in the transition matrix computed by the *teSMC* method. Both methods belong thus to the same conceptual framework and are complementary.

The agreement between our estimations for the transition to selfing in *A. thaliana* and the ones obtained based on *S*-locus analyses is remarkable, because they show that important evolutionary insights can be obtained with small amount of data, when this data is analyzed using appropriate adhoc methods relying on evolutionary theory (Bechsgaard, et al. 2006). However, *S*-locus based inferences of shifts to selfing are limited to species for which such transitions have been caused by a loss-of-function mutation in the *S*-locus and for which the *S*-locus has been identified and properly assembled. This information is only partially available for other plant species, and the genetic determinism of selfing also varies between genera (Franklin-Tong 2008). Thus, our inference methods offer the opportunity to test for the existence and the timing of changes in mode of reproduction in potentiallany species providing full-genome polymorphism data are available. They also pave the way for addressing long standing questions on the evolution of reproductive systems which cannot be directly tested from field experimental or even phylogenetic approaches. How frequent and how recent are transitions from outcrossing to selfing is the main question, and the one that motivated our work. Phylogenetic methods can at best infer that a transition has occurred at some time on a given branch of a phylogeny. For example, considering the case of *A. thaliana*, knowing that the two sister species, *A. lyrata* and *A. haleri*, are self-incompatible indicates that the shift to selfing occurred after the divergence between the two lineages, that is between present time and 13 millions years (Beilstein, et al. 2010), which is poorly informative. In addition, from phylogenetic character mapping, when two or more sister species share the same character state, most of the time the shift to this state is inferred before the divergence of the species. However, it may not be the case for very labile traits with high extinction rate as self-fertilization. In the fungus genus *Neurospora*, although a clade of species shares a homothallic mating system (equivalent to selfing), a detailed molecular analysis of the *mat* locus that controls mating system revealed that the breakdown of this locus occurred several times independently (Gioti, et al. 2012). Reversion from selfing to outcrossing is supposed to be very rare but this question is still debated (Barrett 2013). Our methods provide the adequate tool to tackle this question and their systematic application to various species may help discovering such possible reversion or even more complex scenarios. As an example, we apply *teSMC* to estimate scenarios of transition from outcrossing to selfing followed by reversion, or the reverse, as well as more gradual (stepwise) transitions (Figure S13 & Figure S14). The method performed well and is thus promising for detecting more complex histories.

Another useful application of the methods is for demographic inferences when shifts in mating systems are suspected. As they alter the distribution of age and lengths of *T*_MRCA_-segments they can lead to spurious shifts in population size if not taken into account (Figure S6). For example, using an SMC approach, ancestral drops in population sizes have been inferred in the selfers *Capsella orientalis* and *C. bursa-pastoris* but not in the outcrosser *C. grandiflora* (Kryvokhyzha, et al. 2019). This could correspond to real changes in population size but also to transition towards selfing. This issue is especially worth being considered in cultivated species. The demographic history associated with domestication is a central question for the study of crop species and demographic scenarios have been inferred in many species. However, shifts, or at least variations, in mating system occurred quite frequently during plant domestication (ex: African rice, tomato, grapevine, melon) (Glémin and Bataillon 2009; Meyer, et al. 2012) and taking such variations into account would help refining demographic scenarios.

The methods we developed focus on outcrossing/selfing transitions. However, they could be extended to other reproductive modes such as sex/asex transitions by adapting the relevant population parameters using existing works on coalescence with facultative sexual reproduction (Hartfield, et al. 2018). Overall, our methods may open new ways to answer the old riddle of why so many species do reproduce sexually (Barton and Charlesworth 1998).

## Materials and Methods

### Modelling of a transition to selfing using forward and coalescent simulations

To model a transition from outcrossing to predominant selfing, we considered a single population composed of *N* diploid individuals. At each generation, each offspring is generated by self-fertilization of a single individual or by outcrossing with probabilities σ and 1-σ, respectively, where *σ* is the selfing rate. Unless stated otherwise, transitions to predominant selfing were modelled by allowing the selfing rate to change instantaneously from *σ*_ANC_ to *σ*_PRES_ at time *t_σ_*. The mutation and recombination were set to 1×10^-8^ events per generation per nucleotide. When needed the population size was allowed to change instantaneously from *N*_ANC_ to *N_PRES_* at *t*_N_. This model was implemented using the Wright-Fisher (WF) simulation mode in *slim3* (Haller and Messer 2019); scripts to simulate genetic data using this model are available on our git repository (https://github.com/laurentlab-mpipz/struett_and_sellinger_et_al.git). We used this model to generate genetic variation for a sample of *n* = 20 haploid genomes, sampled from 20 different individuals, and composed of five DNA sequences of 1Mb each.

The same model was implemented in a coalescent framework using *msprime* (Kelleher, et al. 2016). Following Nelson, et al. (2020) who showed that continuous-time coalescent simulations of large sequences cause biases in patterns of identity-by-descent and linkage disequilibrium, we implemented a hybrid model in which the first 1,000 generations were simulated using a discrete-time coalescent process and the following generations were modelled using the SMC’ algorithm (Marjoram and Wall 2006). The coalescent implementation, which runs significantly faster than the forward-WF implementation, was used for the ABC and performance analyses (see below), while the WF-forward implementation was used to assess the quality of the coalescent-with-selfing approximation proposed by (Nordborg and Donnelly 1997; Nordborg 2000) (Figure S1) and was extended to study the consequences of negative selection on inference with the *tsABC* and *teSMC* methods (see below).

### Analysis of T_MRCA_-segments in simulated data

For the forward and coalescent simulations, we wrote functions to obtain the lengths and the time to the most recent ancestor (*T*_MRCA_) of *T*_MRCA_-segments. *T*_MRCA_-segments are sets of contiguous nucleotides in a sample of size two that share the same MRCA. The joint distribution of *T*_MRCA_ and lengths of those segments (TL-distributions) was used to describe the consequences of a transition to selfing at the genomic level and how it differs from a change in population size (Figure 1, Figure S1, Figure S3). *T*_MRCA_-segments were analyzed by identifying consequential recombination events in the history of the sample (i.e. events that lead to the inclusion of a new MRCA) and the corresponding breakpoints represented the boundaries of the successive segments. Then, the *T*_MRCA_ of each segment was obtained by identifying the MRCA of each segment.

We also calculated the transition matrix of *T*_MRCA_ of successive *T*_MRCA_-segments along the genome (TM_true_, Figure 1D,E). For this, we discretized *T*_MRCA_ values and counted the frequencies of segment transitions along simulated sequences for each combination of discrete *T*_MRCA_ values. *T*_MRCA_ were discretized using a similar approach as in MSMC (Schiffels and Durbin 2014) with the lower boundary of bin *i* given by, -8*N* x log(1-*i*/*m*), where *N* is the population size and *m* the total number of bins. The relevant code can be found at https://github.com/laurentlab-mpipz/struett_and_sellinger_et_al.git.

### Calculation of summary statistics of polymorphism data

While TL-distributions carry the characteristic signature of shifts to selfing, they are also challenging to infer from empirical genetic data. Therefore, we used three summarization approaches to capture this signal using polymorphism data: 1) the unfolded site frequency spectrum (SFS), which is the distribution of absolute derived allele frequencies in the sample and is known to carry information about past population size changes. 2) A discretized distribution of linkage disequilibrium (LD) decay inspired from the approach taken by Boitard, et al. (2016) who used it jointly with the SFS to estimate past changes in population sizes. Unlike the SFS, which only carries information about *N* but not the recombination rate (*r*), LD-decay depends on the product of *N* and *r*. Combining both distributions therefore allows to capture the signature of changes in *N* and *r*. LD was calculated as *r*^2^ from a subset of 10,000 randomly chosen SNPs and discretized into discrete physical distances with following breakpoints: 6,105; 11,379; 21,209; 39,531; 73,680; 137,328; 255,958; 477,066; 889,175 bp. 3) Window-based transition matrix (TM_win_): While TM_true_ carries a characteristic signal to estimate shifts to selfing, it is not straightforward to calculate it using polymorphism data. This is because the boundaries of *T*_MRCA_-segments are not directly observable and need to be inferred themselves. TM_win_ captures some of the information in TM_true_ by computing the pairwise diversity in non-overlapping successive windows of 10kb for a sample of size two.

### Simulations with background selection (BGS)

Simulations with BGS were conducted with *slim3*. We used the distribution of fitness effects (DFE) estimated by *DFEalpha* for *A. thaliana* published by Hamala and Tiffin (2020). The DFE was used to assign negative selection coefficients to simulated coding non-synonymous genetic variants only (i.e. we did not simulate negative selection on functional non-coding regions). We took care of simulating realistic proportions and spatial distributions of coding sequences by using the positional information of CDS from the annotation of the reference genome of *A. thaliana* (Arabidopsis Genome 2000). Except for the DFEs and genetic structure all other parameters and dataset dimensions were identical to the simulations without negative selection.

### Application to Arabidopsis thaliana

To obtain the observed summary statistics, we used the imputed genotype matrix provided on the 1001 genomes website (https://1001genomes.org/data/GMI-MPI/releases/v3.1/SNP_matrix_imputed_hdf5/). We used samples from three separate genetic clusters: central European CEU, Iberian non-relicts (IBnr), and Relicts. Only accessions with cluster membership > 95% were included. The final sample sizes are provided in Table 1. We included five genomic regions from the five chromosomes (Table S4). The regions were chosen based on homogeneity of recombination rates and diversity (Figure S15). We resampled 12 haplotypes multiple times and calculated the combined summary statistics, SFS, LD and TMwin. We centralized and normalized the statistics and calculated the first 20-PLS (see Appendix B).

*tsABC* was conducted on the A. thaliana data, by using 12 samples for 5 independent loci of 1 Mb length. The mutation and recombination rates were set to 6.95e-9 (Ossowski, et al. 2010) and 3.6e-9. The recombination rate is the genome-wide average provided by Salomé, et al. (2012). We simulated a total set of 130,000 vectors of summary statistics. Parameter estimates were conducted as described for the ABC performance analysis. We provided the mode of the average posterior distributions as a final result. In addition, we used a subsample of 20 sequences of the *Arabidopsis* data per genetic cluster to estimate demography and the transition to selfing using *teSMC* in the one-transition mode. Data and scripts can be found at (https://github.com/laurentlab-mpipz/struett_and_sellinger_et_al.git) and (https://github.com/TPPSellinger/eSMC2).

## Data availability

A complete detailed description of the *teSMC* and *tsABC* methods is in the SI text appendix A and B. Scripts for all figures, simulations, and the *tsABC* workflow can be found on github (https://github.com/laurentlab-mpipz/struett_and_sellinger_et_al.git). The simulator used to simulate selfing transition as well as *teSMC* are available on github (https://github.com/TPPSellinger/eSMC2). A tutorial is also provided to simulate and analyze data.

## Acknowledgements

This work was funded by the Deutsche Forschungsgemeinschaft (DFG, German Research Foundation), project 462181533 to SL and projects 317616126; 254587930 to AT.

---

**Figure S1.**
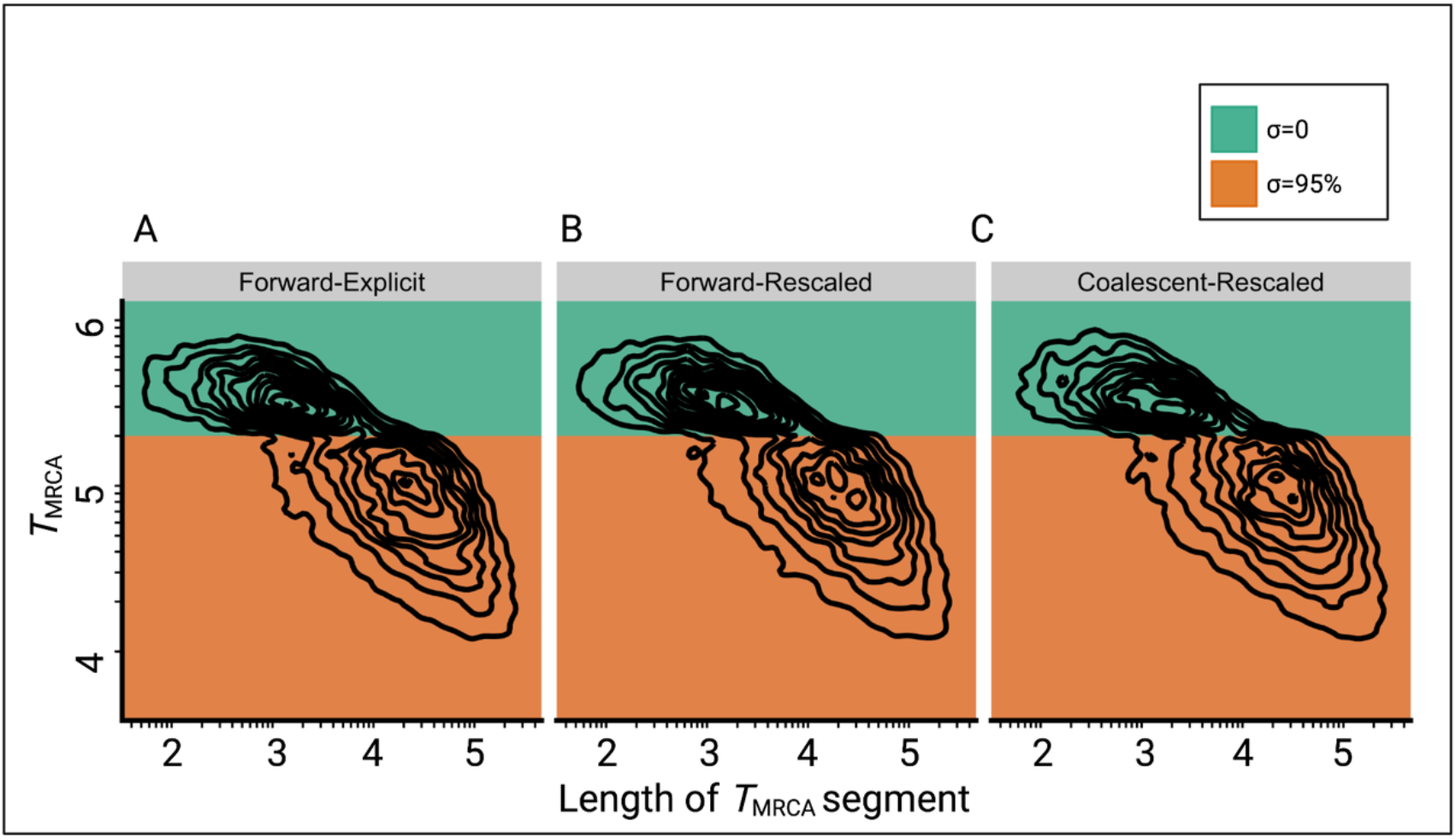
Comparison of the joint distributions of *T*_MRCA_ and lengths of *T*_MRCA_-segments (TL-distribution) under three different simulation approaches. **A:** Explicit selfing implemented in a forward-in-time Wright-Fisher model (*slim3*). Population size is constant and selfing rate shifts from outcrossing (0 = 0, green) to predominant selfing (0 = 0.95, orange). *T*_MRCA_-segments were defined as contiguous sets of nucleotides sharing the same *T*_MRCA_. **B:** Shift to selfing is simulated using a forward-in-time Wright-Fisher model (*slim3*) by rescaling population size and recombination rate at *t*_α_ as suggested by Nordborg and Donnelly (1997) and Nordborg (2000) **C:** Shift to selfing simulated using the coalescent (*msprime*) by rescaling population size and recombination rate at *t*_α_ as in panel B. Axes dimensions are on a log10 scale.

**Figure S2.**
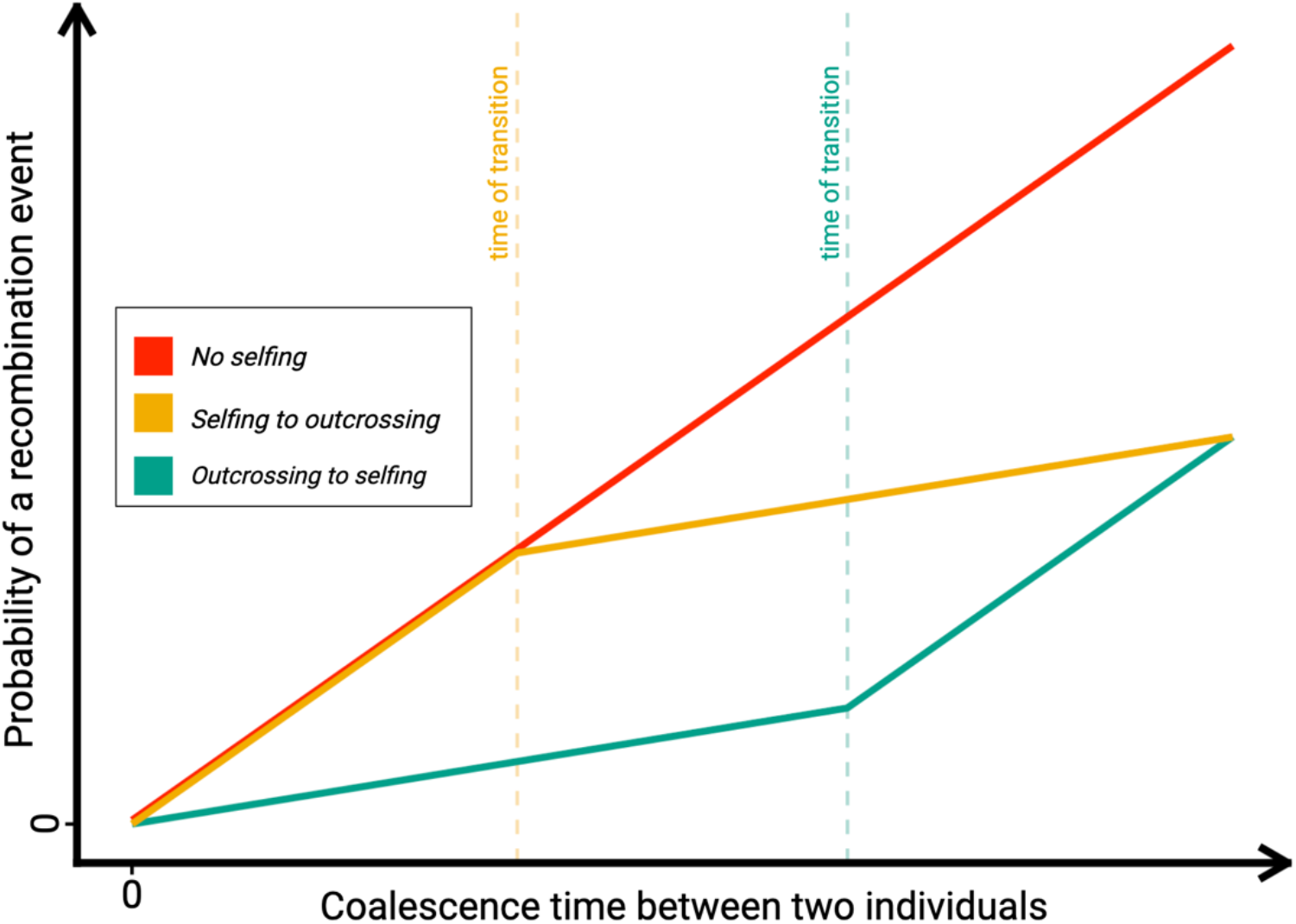
Probability of recombination events over time under different reproductive histories. Schematic representation of the probability of a recombination event to occur in a coalescent tree of sample size two in the absence of selfing (red); in the case of a transition from selfing to outcrossing (forward-in-time, green), and in the case of a transition from outcrossing to selfing (forward-in-time, yellow). The probability of a recombination event having occurred depends on the effective recombination rate, which is in turn negatively impacted by selfing rate.

**Figure S3.**
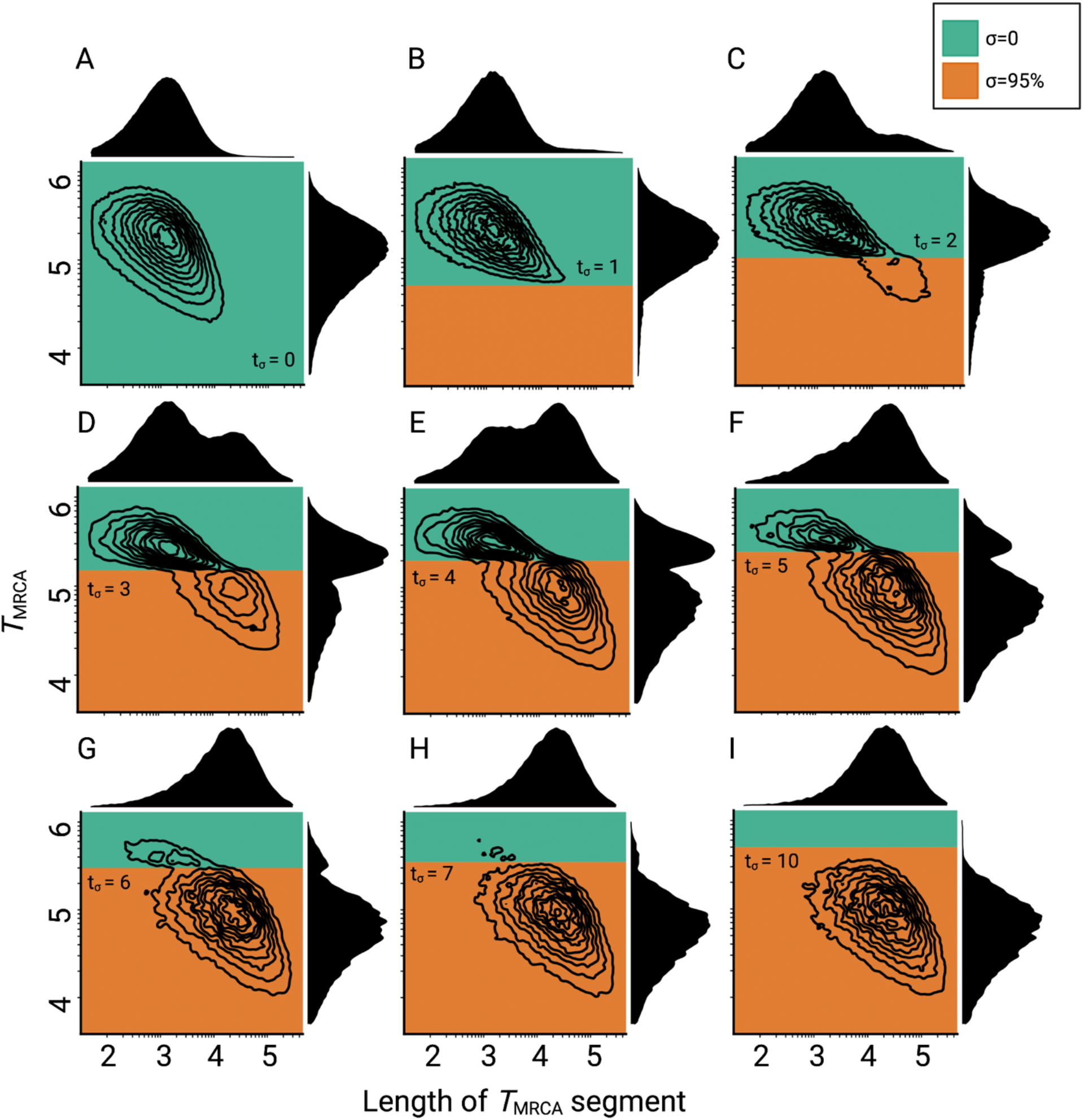
Consequences of a transition to selfing on the genealogies of simulated chromosomes for different ages of the transition. **A-I:** Joint and marginal distributions of ages in generations and lengths of *T*_MRCA_-segments in a population with a constant population size and a shift from outcrossing (green) to predominant selfing (orange). *T*_MRCA_-segments were defined as contiguous sets of nucleotides sharing the same most recent common ancestor. Transitioning times are provided on *N*σ scale. Axes dimensions are in log10.

**Figure S4.**
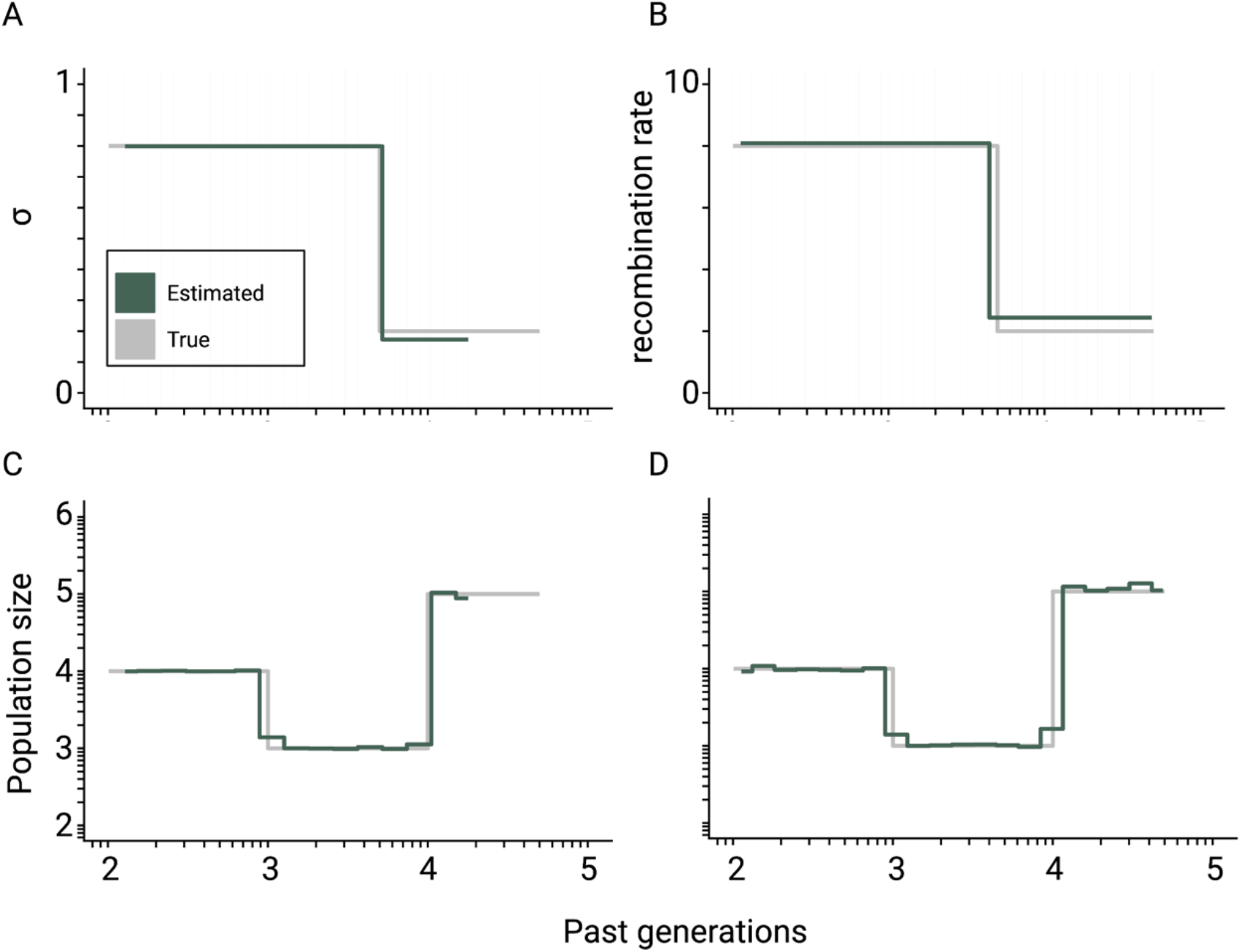
Theoretical convergence of *teSMC* under complex demography. Best-case convergence of *teSMC* using 10 sequences (i.e. haploid genomes) of 100 Mb (green) when population undergoes a bottleneck (true sizes are indicated in grey) with either variation of selfing in time (A,C) or variation of recombination rate in time (B,D). The selfing rate through time is represented in A) and the corresponding estimated population size is represented in C); the recombination rate was set at 1×10^-7^ per generation per bp. The estimated recombination rate trough time is represented in B), it changes from 2×10^-8^ to 8×10^-8^ per generation per bp, and the corresponding population size within D). The mutation rate is set in all cases to 1×10^-8^ per generation per bp.

**Figure S5.**
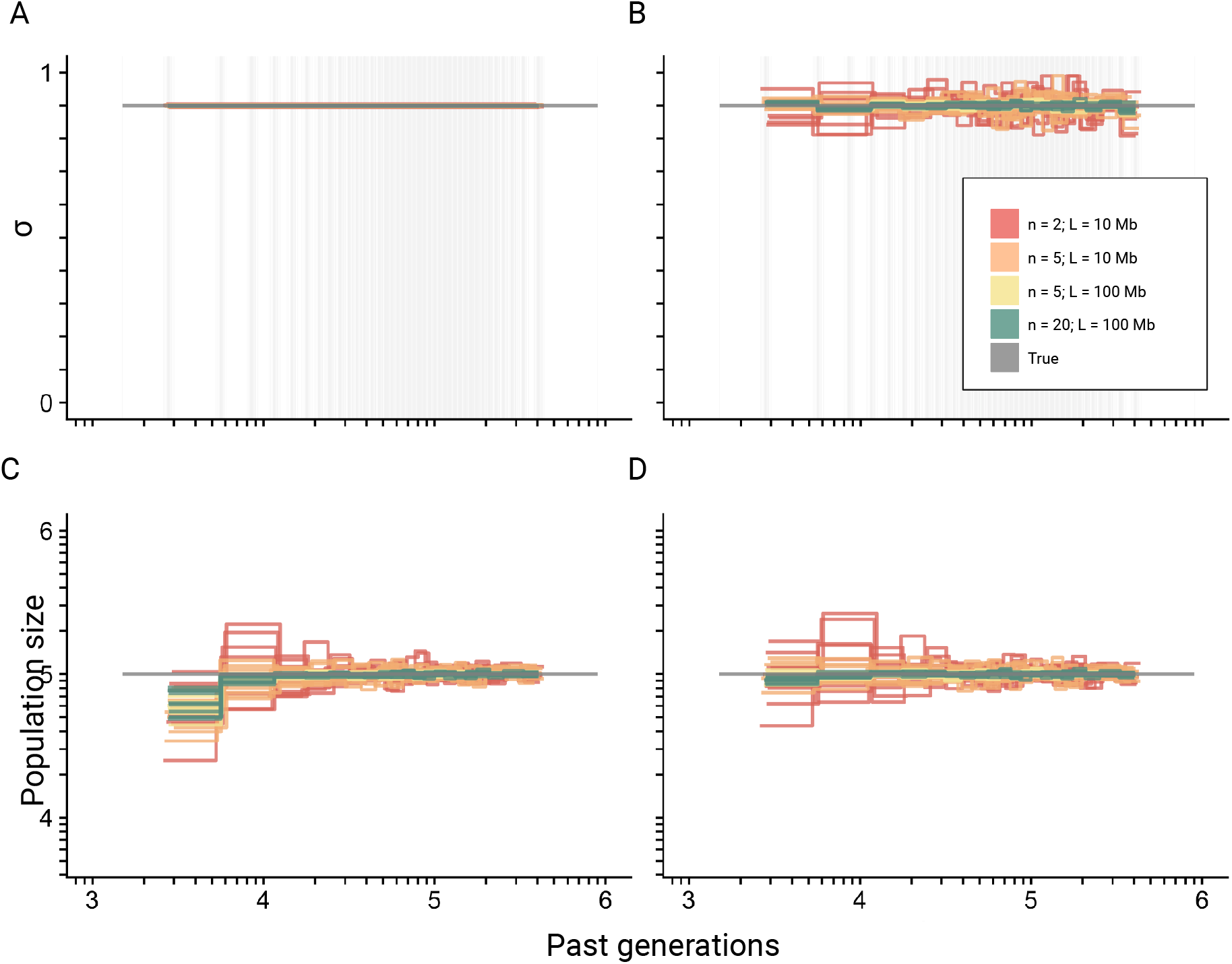
Best-case convergence of *teSMC* for different amount of data. Best-case convergence of *teSMC* using different combinations of sample sizes (*n* = 2, *n* = 5, or *n* = 20 sequences; i.e., haploid genomes) and sequence lengths (L =10Mb or L=100Mb, see legend) Population size is constant (N=100,000, grey horizontal line) with a constant selfing rate of 0.9. The best-case convergence is estimated assuming that selfing is constant (A, C) or varying in time (B, D). The estimated population size assuming constant selfing in time is represented in (C) and the simultaneously estimated selfing rate in (A). The estimated population size assuming varying selfing rate in time is represented in (D) and the simultaneously estimated selfing rate through time in (B). The recombination was set to r=1×10^-8^ per generation per bp. Except for the selfing rates, both axes are on a log10 scale.

**Figure S6.**
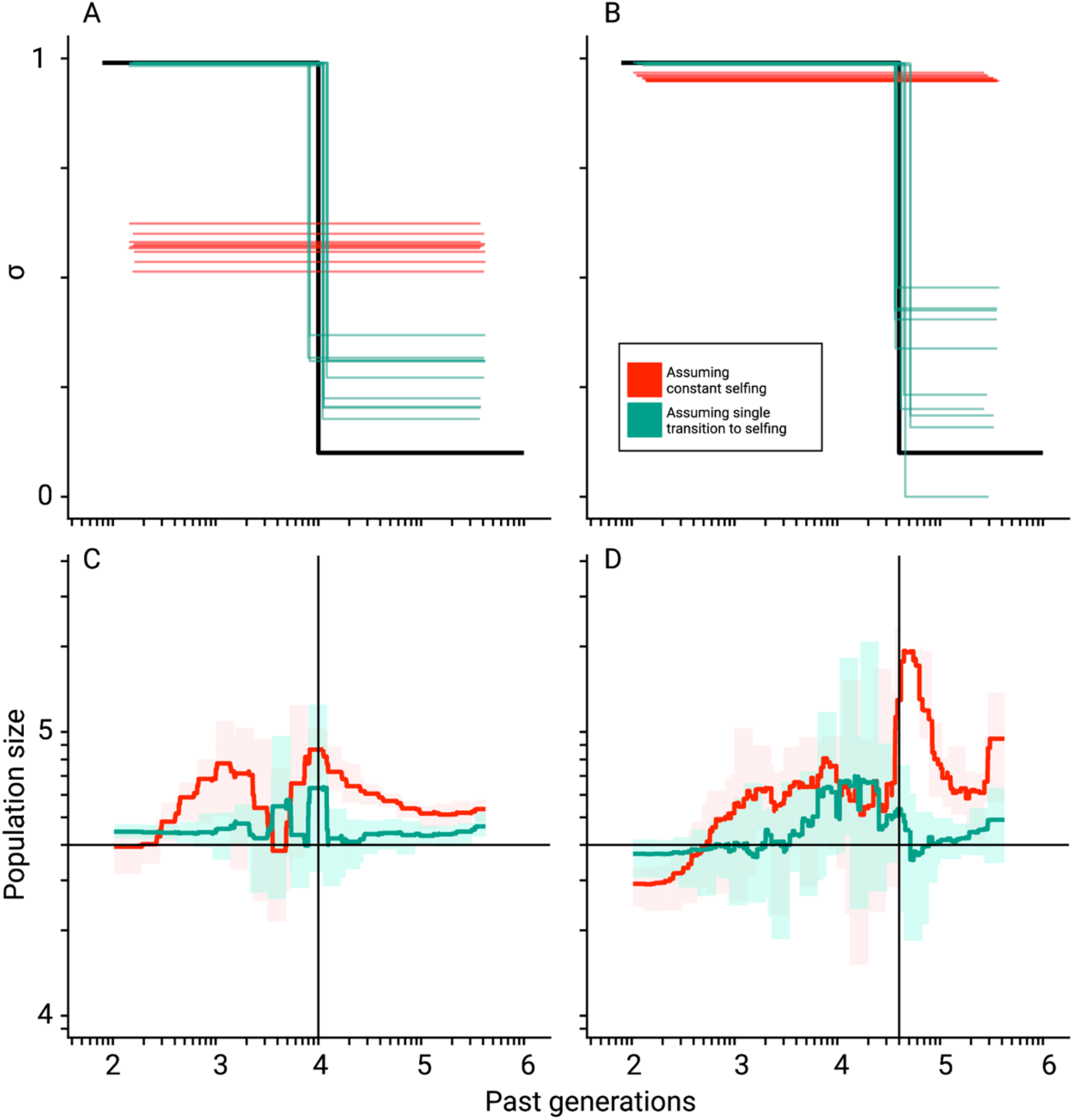
Misinference of population sizes when transitions to selfing are not accounted for. Comparisons between true (black lines) and estimated selfing rates and population sizes estimated by *teSMC* for 10 replicates. Here simulations were done using a constant population size (N=40,000) and a transition to selfing from σ_ANC_ = 0.1 to σ_PRES_ = 0.99 at *t*_σ_ = 10,000 (A,C) and 40,000 (B,D). Five chromosomes of 1Mb each were simulated with mutation and recombination rates set to 1×10^-8^ events per generation per bp. Red and green lines indicate results obtained assuming the wrong model (i.e. constant selfing) and the correct model (i.e. single-transition), respectively. For the selfing rates (A,B), results for each replicate are indicated with solid lines. For the population sizes, the 10 replicates were summarized by the green and red shaded areas, where the width of the shaded area corresponds to the range between the minimum and maximum value observed across replicates.

**Figure S7.**
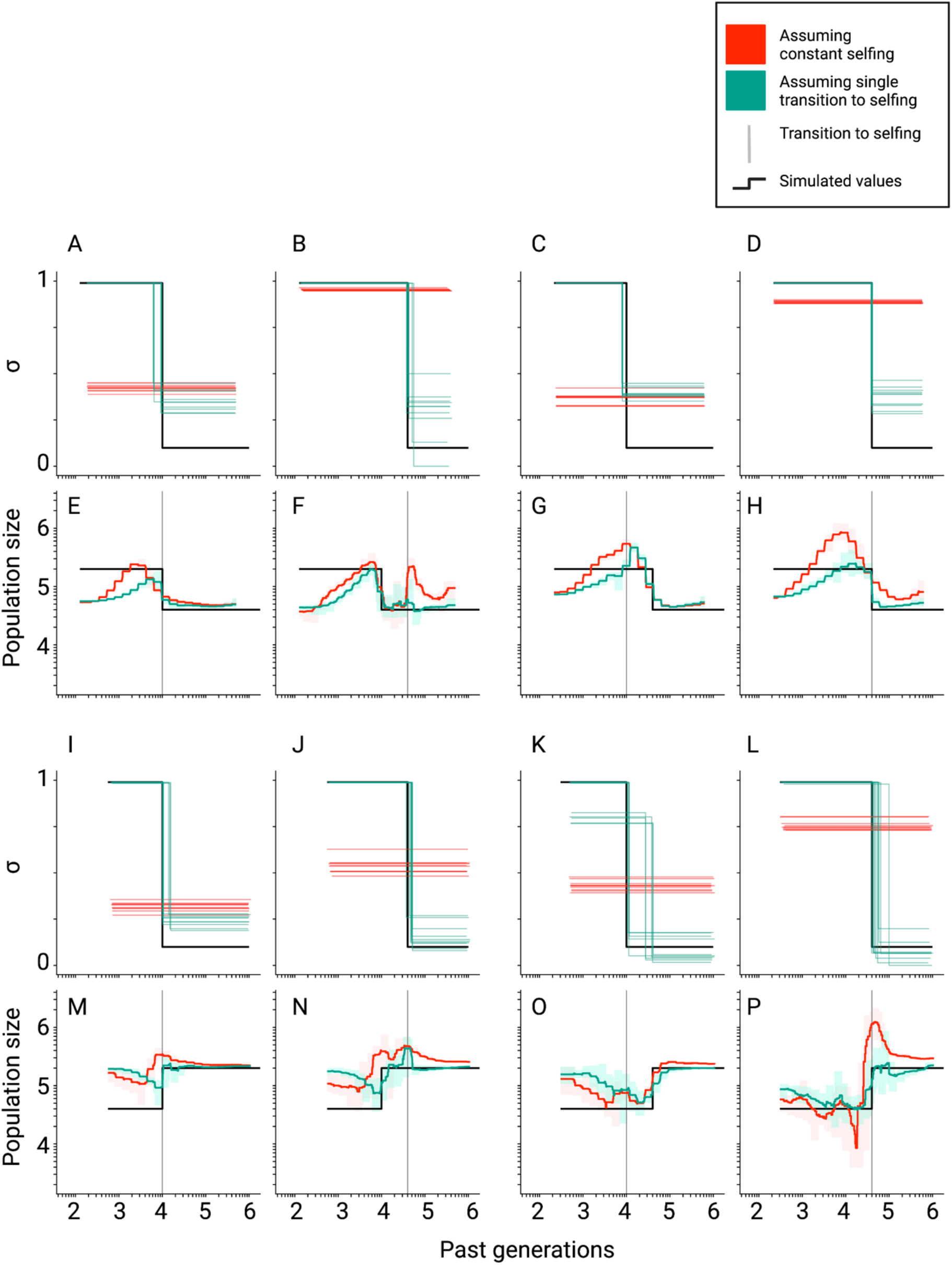
Inference of population sizes and selfing rates estimated by *teSMC* when both parameters change over time. **A – P**: Comparisons between true (black lines) and estimated selfing rates and population sizes estimated by *teSMC* for 10 replicates. Here simulations were done as in Figure S6 except for the addition of a single stepwise population size expansion forward-in-time (first and second rows) or contraction (third and fourth row). The transition to selfing occurred from σ_ANC_ = 0.1 to σ_PRES_ = 0.99 at *t*_σ_ = 10,000 (A, C, E, G, I, K, M, O; first and third column) and 40,000 (B, D, F, H, J, L, N, P; second and forth column). Red and green lines indicate results obtained assuming the wrong model (i.e. constant selfing) and the correct model (i.e. single-transition), respectively. For the selfing rates results for each replicate are indicated with solid lines. For the population sizes, the 10 replicates were summarized by the green and red shaded areas, where the width of the shaded area corresponds to the range between the minimum and maximum value observed across replicates.

**Figure S8.**
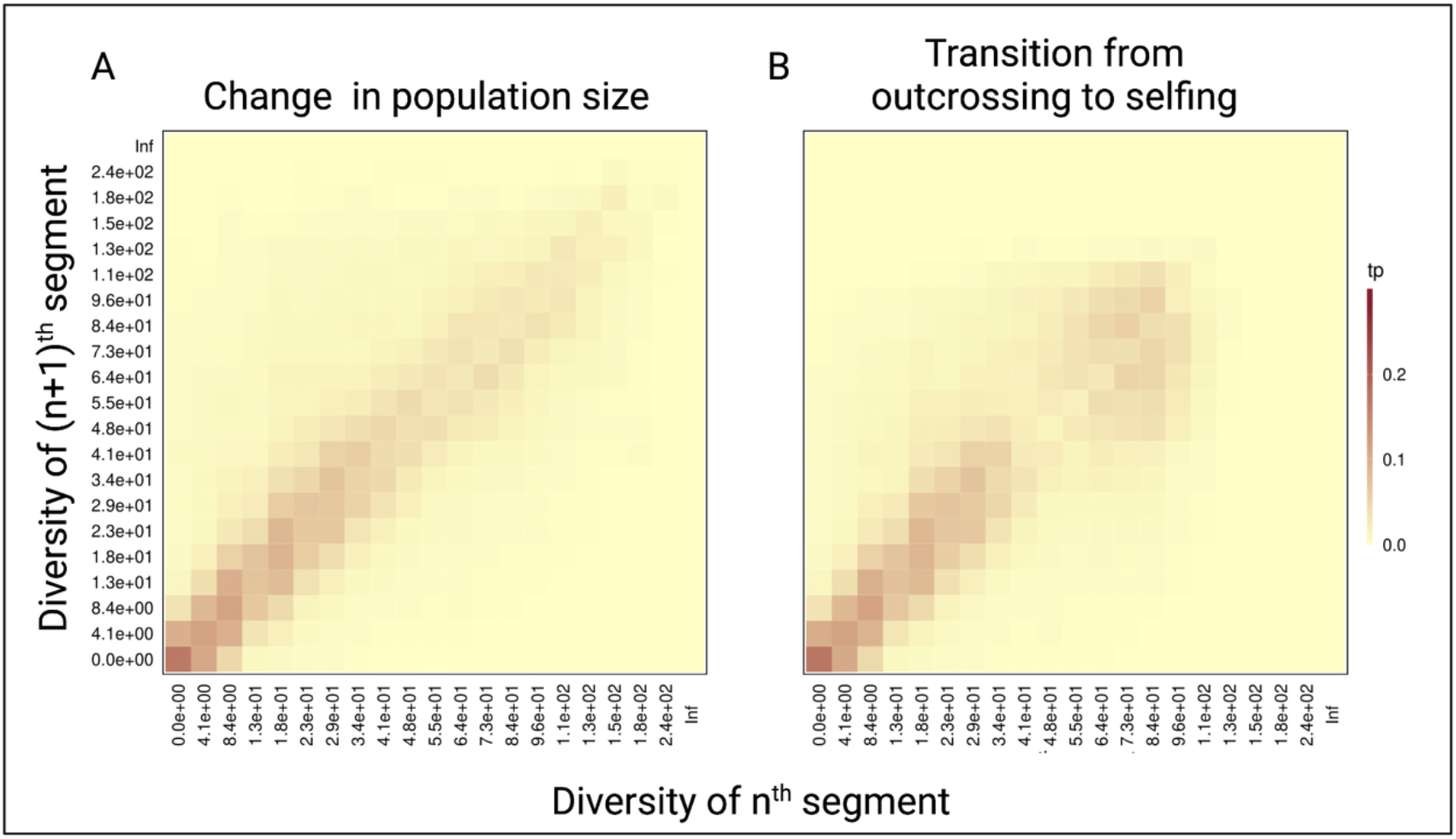
Pairwise diversity transition matrices used in *tsABC*. **A:** The transition matrix of pairwise diversity (TMwin) between adjacent non-overlapping 10kb windows measured for 1Mb of data simulated with the population-size-change-model (Figure 3B) in an outcrossing population with a stepwise change from *N*_ANC_ = 100,000 to low *N*_PRES_=50,500. The population sizes were chosen to correspond to the rescaling of the effective population size by the selfing rates used in panel B. **B:** The same transition matrix of pairwise diversity as in panel A for a constant population (*N*=100,000) but with a transition from outcrossing to predominant selfing (*σ*=0.99). The recombination rate was set to 1×10^-8^. These matrices summarize the probabilities that the *n*^th^ window with a given diversity *X* is followed by the (*n*+1)^th^ window of diversity *Y*. The heat colors indicate the transition probabilities (tp).

**Figure S9.**
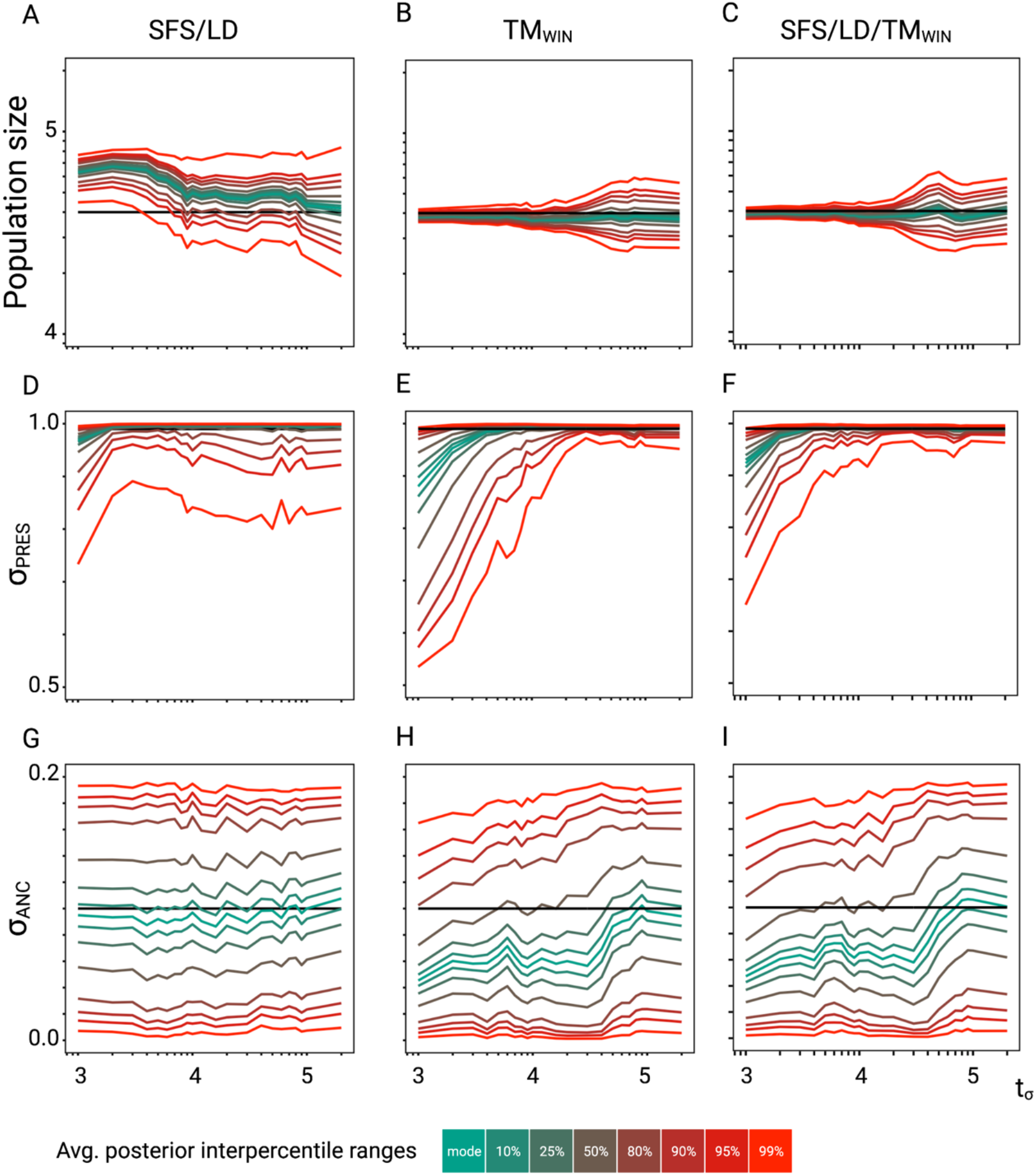
ABC performance analysis. Parameter re-estimation of the three additional parameters of the model described in Figure 3. **A, B, C:** Re-estimation of the population size on 100 datasets simulated under a model with constant population size (N = 40,000) and a change in selfing rate from σANC = 0.1 to σPRES = 0.99. Coloured lines represent the average quantiles for 100 posterior distributions corresponding to the given credible intervals. **D, E, F:** Re-estimation of the present selfing rate on 100 datasets simulated under a model with constant population size (N = 40,000) and a change in selfing rate from σANC = 0.1 to σPRES = 0.99. Coloured lines represent the average quantiles for 100 posterior distributions corresponding to the given credible intervals. **G, H, I:** Re-estimation of the ancestral selfing rate on 100 datasets simulated under a model with constant population size (N = 40,000) and a change in selfing rate from σANC = 0.1 to σPRES = 0.99. Coloured lines represent the average interpercentile ranges for 100 posterior distributions corresponding to the given credible intervals.

**Figure S10.**
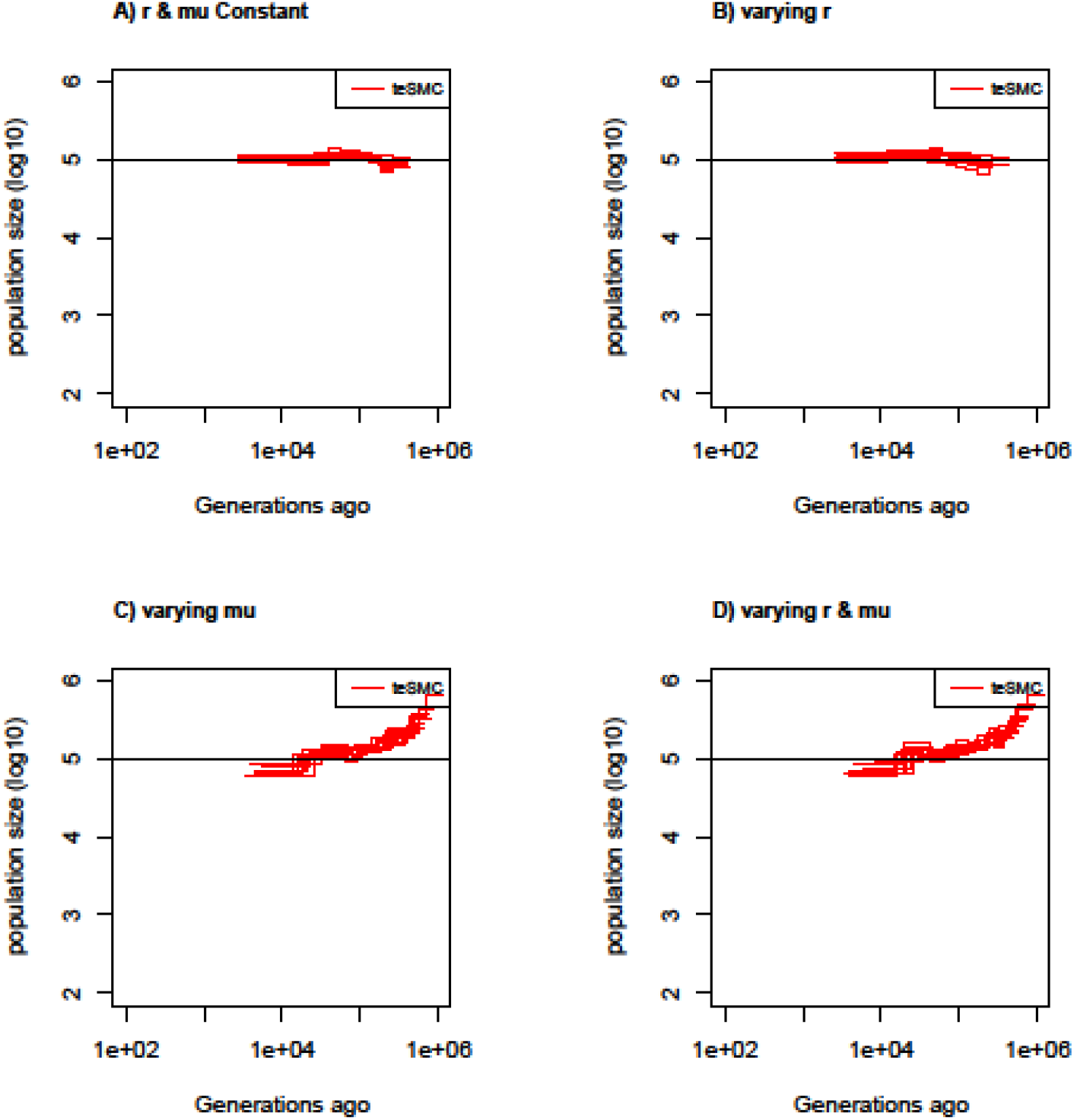
Estimated population sizes by *teSMC* with variable mutation and recombination rates along the genome. Estimated population sizes by *teSMC* using 20 sequences (i.e., haploid genomes) of length 5 Mb (red) when population size is constant and set to 100,000 (black line) with a constant selfing value of 0.9. Simulations are performed under four different conditions (A,B,C,D). A) Recombination and mutation rates are constant along the genome, B) Recombination rate varies randomly between 5×10^-9^ and 2×10^-8^ every 50kb but the mutation rate remains constant. C) Mutation rate varies between 5×10^-9^ and 2×10^-8^ every 50kb but the recombination rate remains constant. D) Mutation and recombination rates both vary along the genome. When constant, recombination and mutation rates are set to 1×10^-8^ per generation per bp.

**Figure S11.**
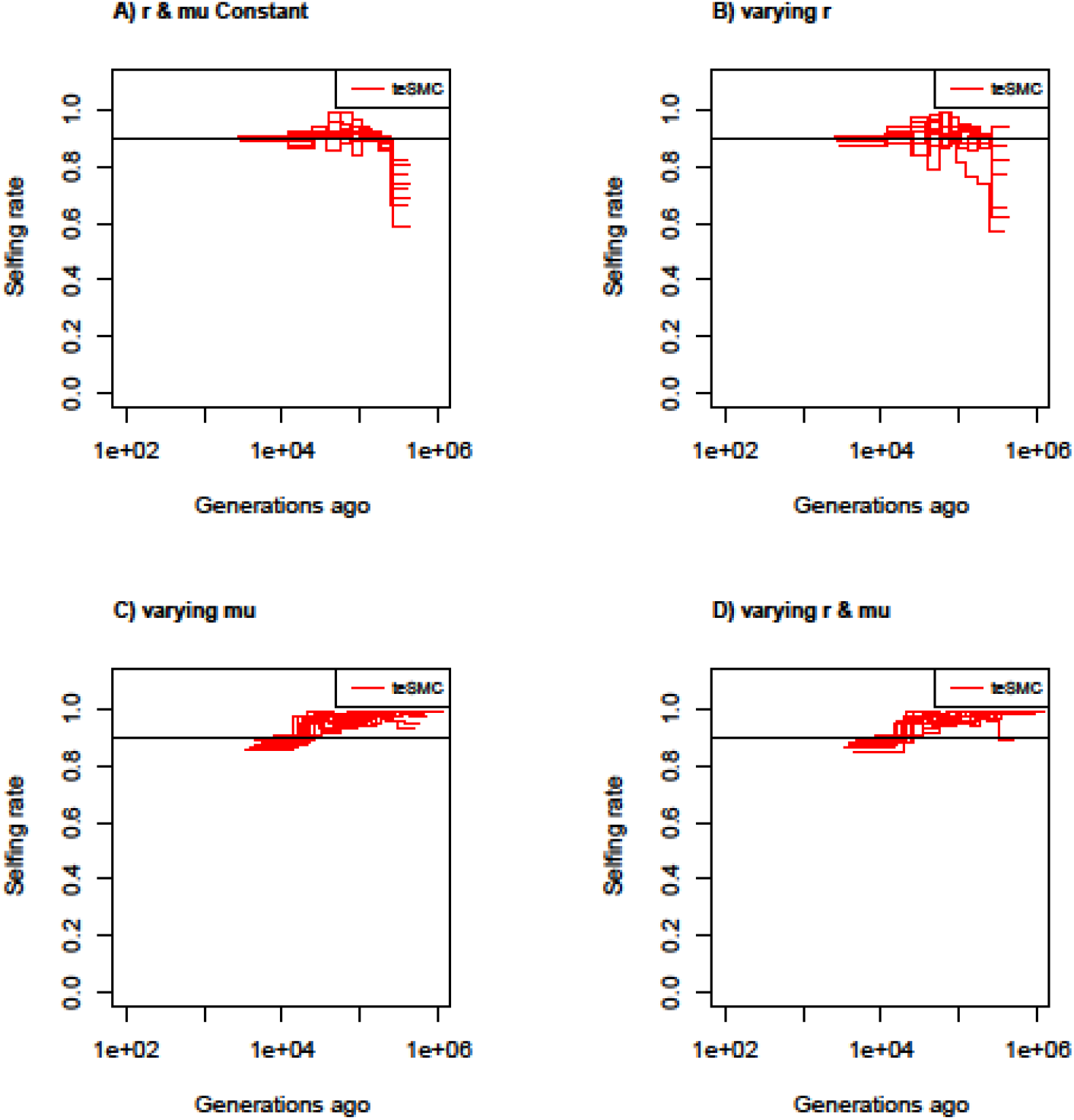
Estimated selfing rates through time by *teSMC* with variable mutation and recombination rates along the genome. Estimated selfing rates through time estimated by *teSMC* using 20 sequences (i.e., haploid genomes) of length 5 Mb (red) when population size is constant and set to 100,000 (black line) with a constant selfing value of 0.9. Simulations are performed under four different conditions (A,B,C,D). A) Recombination and mutation rates are constant along the genome, B) Recombination rate varies randomly between 5×10^-9^ and 2×10^-8^ every 50kb but the mutation rate remains constant. C) Mutation rate varies between 5×10^-9^ and 2×10^-8^ every 50kb, but the recombination rate remains constant. D) Mutation and recombination rates both vary along the genome. The constant recombination and mutation rate are set to 1×10^-8^ per generation per bp.

**Figure S12.**
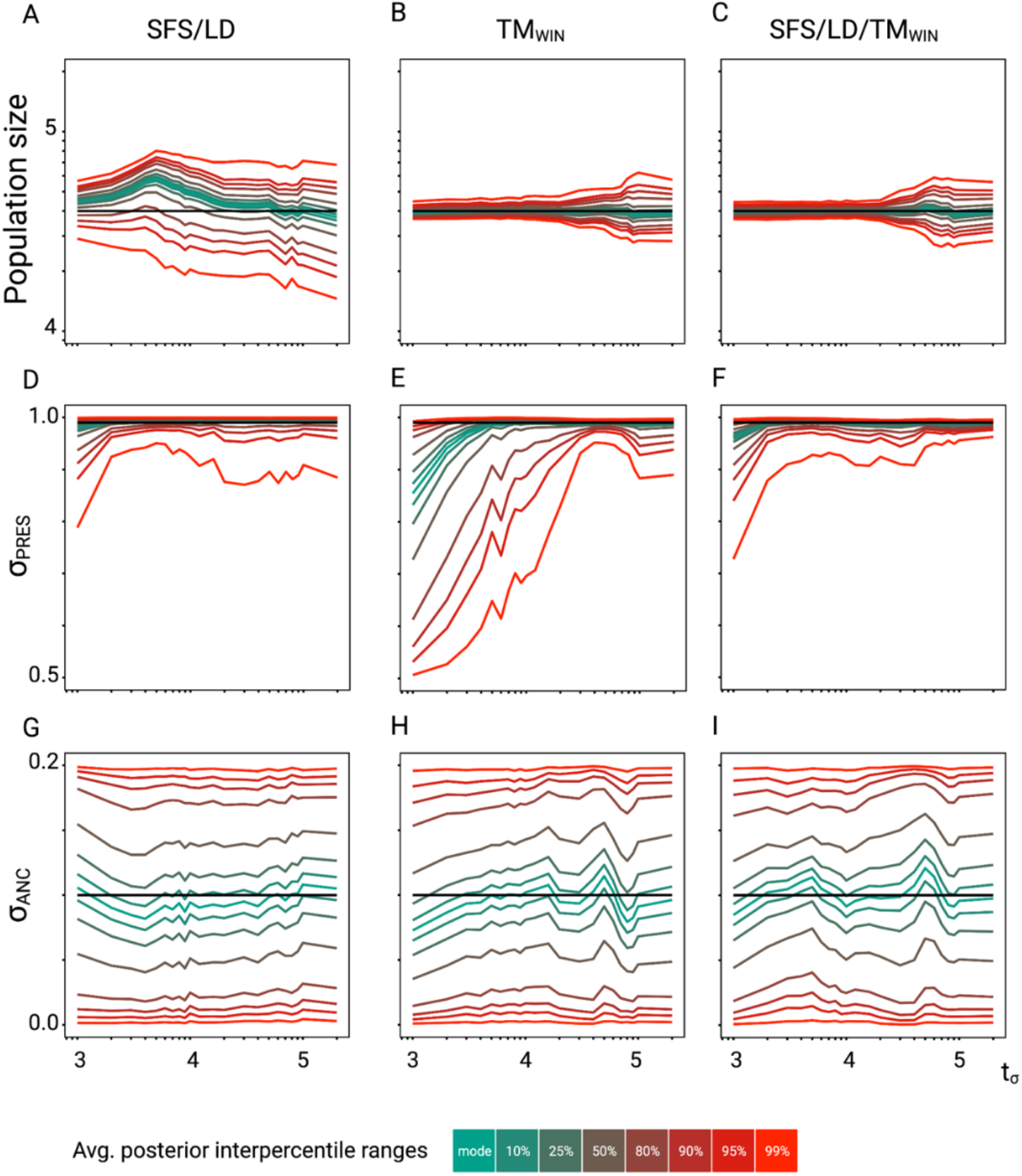
Accuracy of *tsABC* in the presence of background selection (BGS). Parameter re-estimation of the three remaining parameters of the model (see Figure 3A). **A, B, C:** Re-estimation of the population size on 100 datasets simulated under a model with constant population size (N = 40,000) and a change in selfing rate from σANC = 0.1 to σPRES = 0.99. Coloured lines represent the average quantiles for 100 posterior distributions corresponding to the given credible intervals. **D, E, F:** Re-estimation of the present selfing rate on 100 datasets simulated under a model with constant population size (N = 40,000) and a change in selfing rate from σANC = 0.1 to σPRES = 0.99. Coloured lines represent the average quantiles for 100 posterior distributions corresponding to the given credible intervals. **G, H, I:** Re-estimation of the ancestral selfing rate on 100 datasets simulated under a model with constant population size (N = 40,000) and a change in selfing rate from σANC = 0.1 to σPRES = 0.99. Coloured lines represent the average quantiles for 100 posterior distributions corresponding to the given credible intervals. The panels should be compared with the case without linked negative selection in Figure S11. Coloured lines represent the average interpercentile ranges for 100 posterior distributions obtained with *tsABC*.

**Figure S13:**
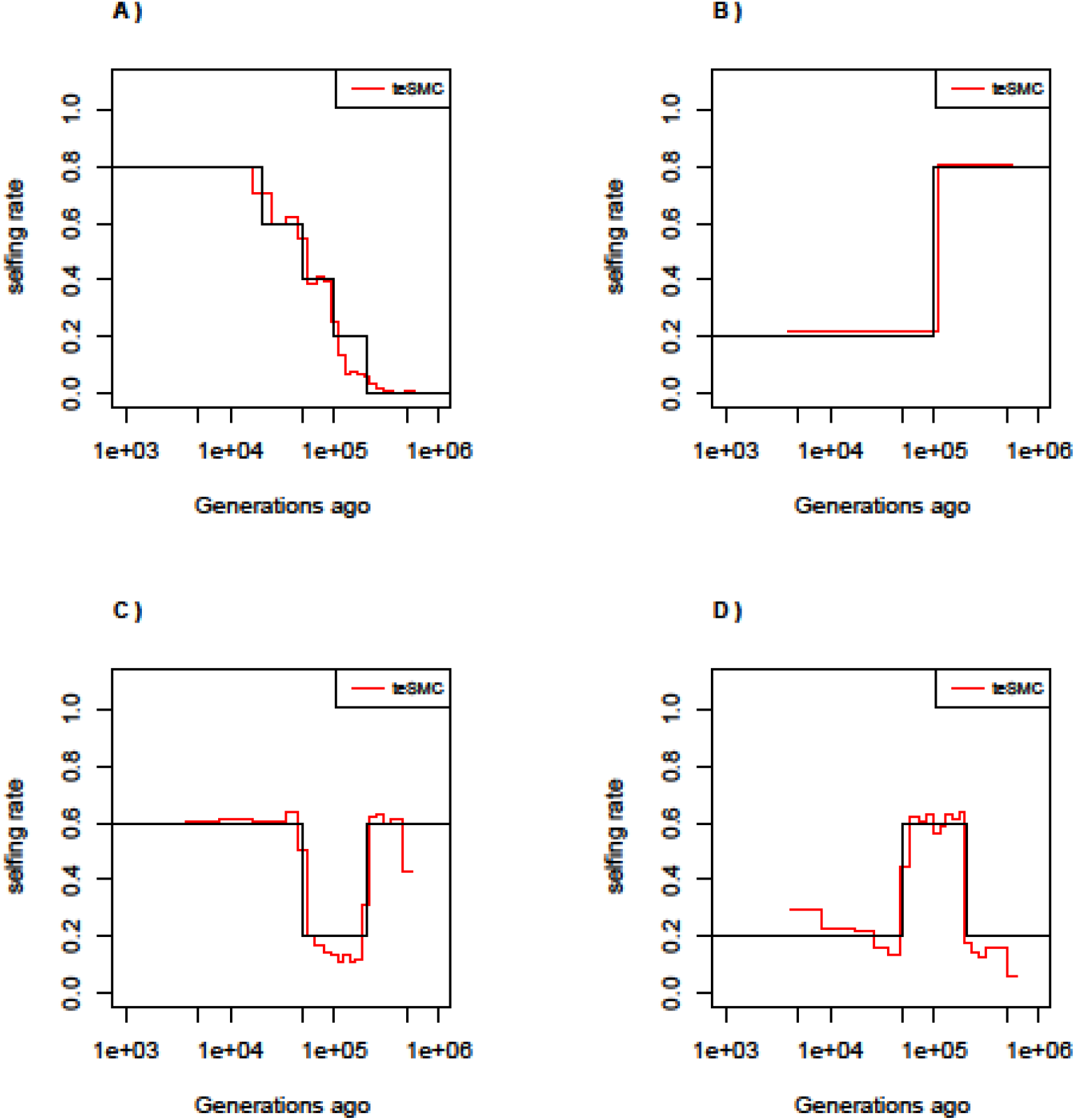
Best-case convergence of *teSMC* under complex selfing transitions. Best-case convergence of *teSMC* under four different scenarios of selfing transition. A) Slow transition from outcrossing to selfing. B) Transition from selfing to outcrossing. C) Transition from 60% selfing to 80% outcrossing to 60% selfing. D) Transition from 20% selfing to 60% selfing to 20% selfing. Population size is set to 100,000 and constant in time. Sample size is set to 20 haploid genome and sequence length to 100 Mbp. Recombination and mutation rate are set to 10^-8^ per generation per bp.

**Figure S14:**
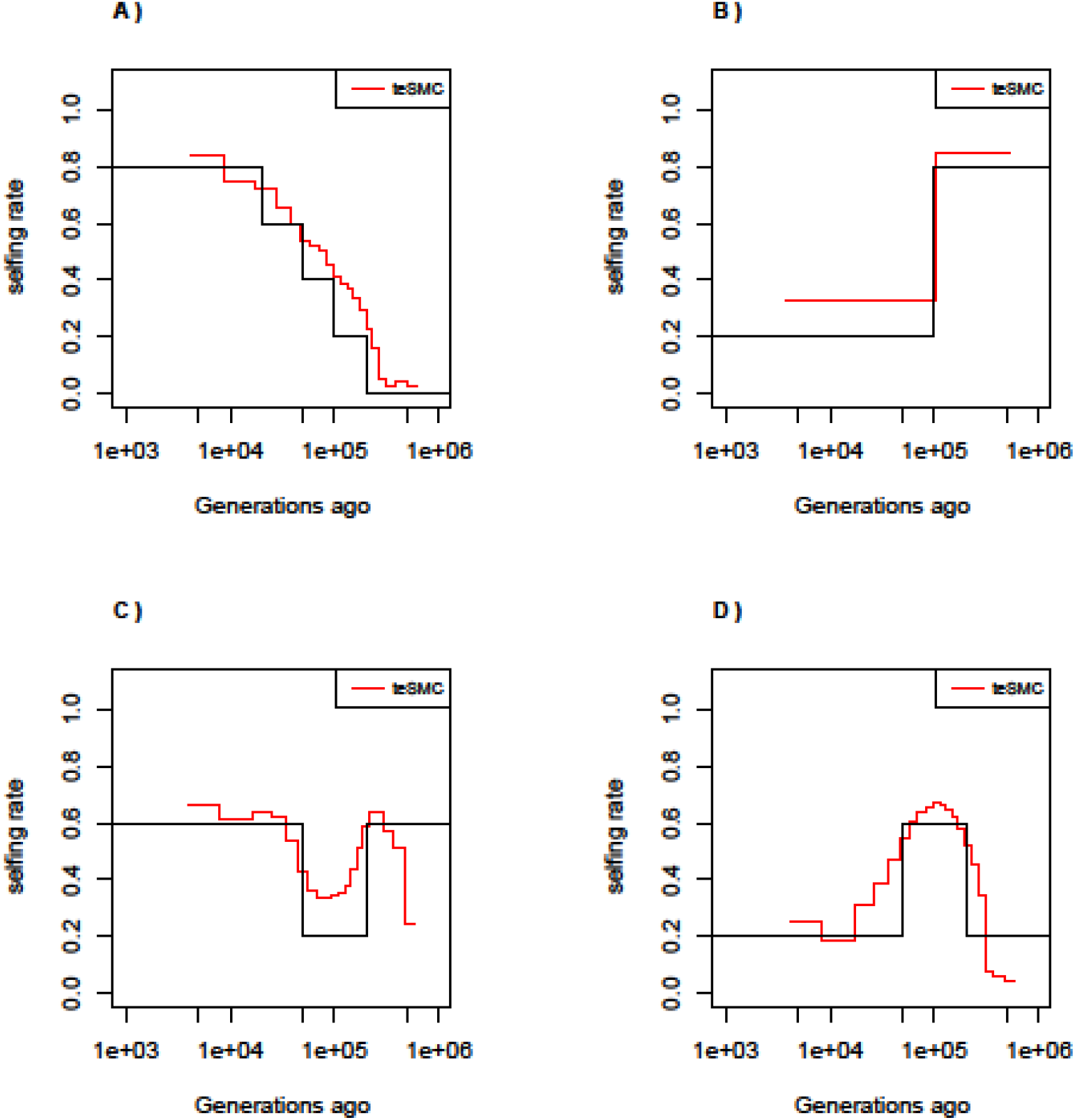
Performance of *teSMC* under complex selfing transitions. Performance of *teSMC* on simulated sequence data under four different scenarios of selfing variations. A) Slow transition from outcrossing to selfing. B) Transition from selfing to outcrossing. C) Transition from 60% selfing to 80% outcrossing to 60% selfing. D) Transition from 20% selfing to 60% selfing to 20% selfing. Population size is set to 100,000 and constant in time. Sample size is set to 10 haploid genomes, with sequence length of 10 Mbp. Mutation and recombination rate are respectively set to 5×10^-8^ and 10^-8^ per generation per bp.

**Figure S15:**
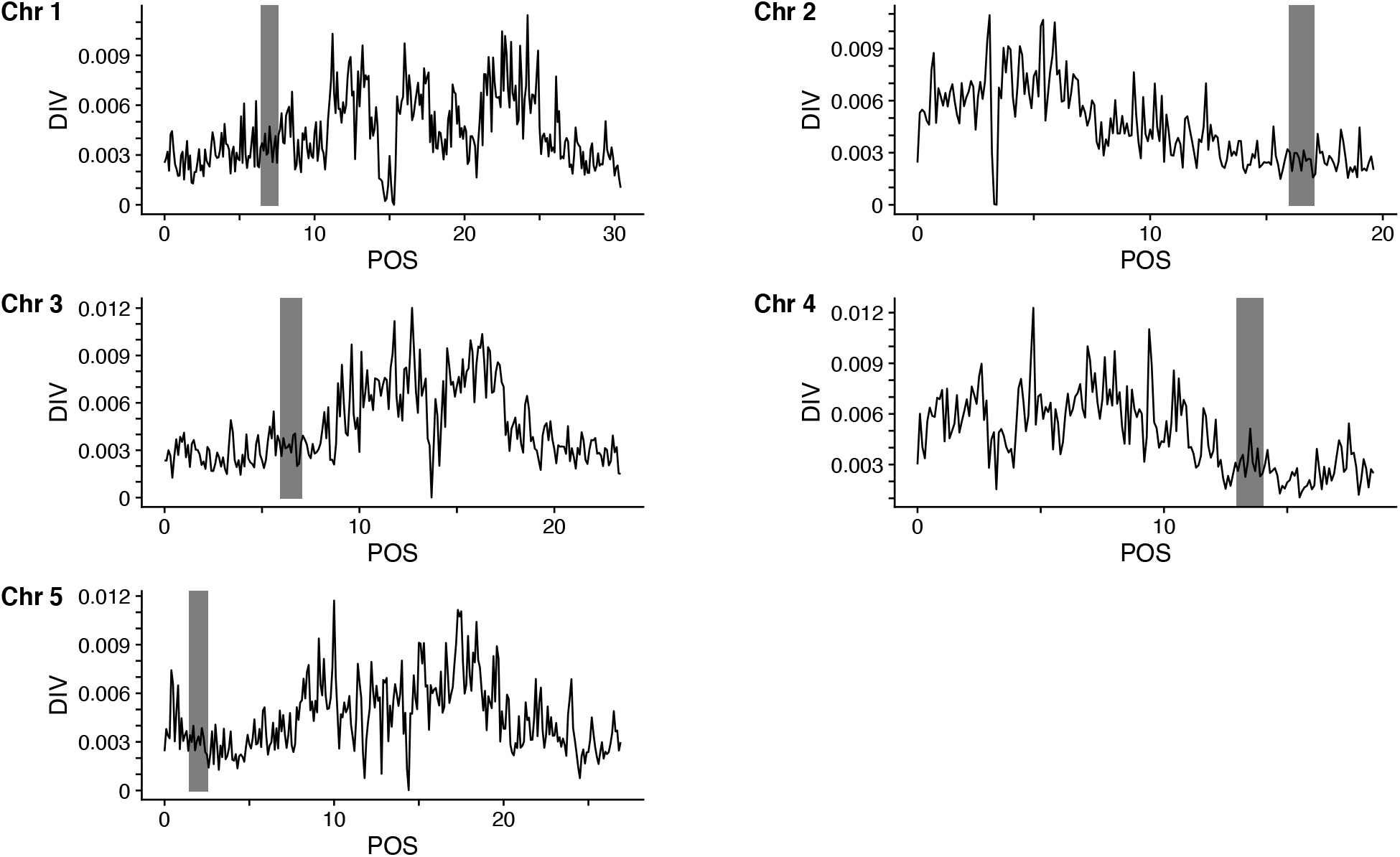
Genomic regions of *A. thaliana* genome (TAIR10) used for the *teSMC* and *tsABC* analyses. Panels show, for the five chromosomes of *A. thaliana,* the nucleotide diversity (pi) calculated with *tskit* using a sliding window of 100 kb. Results are shown only for the CEU sample used for the *teSMC* and *tsABC* analyses. Genomic regions used for the *teSMC* and *tsABC* analyses are highlighted in gray. The same regions were used to obtain the exonic distribution for the BGS simulations in Figure 4.

**Table S1:** Parameters for simulated datasets to investigate the performance of *tsABC*.

| *Model* | *Parameter* | *Value* |
| --- | --- | --- |
| A | N | 40,000.00 |
| A | SIGMA_PRES | 0.99 |
| A | SIMGA_ANC | 0.10 |
| A | T_SIGMA | $t \in (0.05, 10)$ |

**Table S2:** Parameter priors used for the performance analysis of *tsABC*.

| *Model* | *Parameter* | *lower* | *upper* | *type* |
| --- | --- | --- | --- | --- |
| A | N | 10,000.00 | 200,000.00 | logunif |
| A | SIGMA_PRES | 0.50 | 1.00 | uniform |
| A | SIMGA_ANC | 0.00 | 0.20 | uniform |
| A | T_SIGMA | 1,000.00 | 500,000.00 | logunif |
| B | N_ANC | 1,000.00 | 200,000.00 | logunif |
| B | N_PRES | 1,000.00 | 200,000.00 | logunif |
| B | SIGMA | 0.50 | 1.00 | uniform |
| B | T_N | 1,000.00 | 500,000.00 | logunif |

**Table S3:** Parameter priors used to estimate the transition from outcrossing to selfing in *A. thaliana*.

| *Parameter* | *lower* | *upper* | *type* |
| --- | --- | --- | --- |
| N_PRES | 50,000.00 | 500,000.00 | logunif |
| N_ANC | 50,000.00 | 1,000,000.00 | logunif |
| T_N | 10,000.00 | 1,000,000.00 | uni-<br>form |
| SIGMA_PRES | 0.50 | 0.10 | uni-<br>form |
| SIMGA_ANC | 0.00 | 0.20 | uni-<br>form |
| T_SIGMA | 10,000.00 | 500,000.00 | logunif |

**Table S4:**
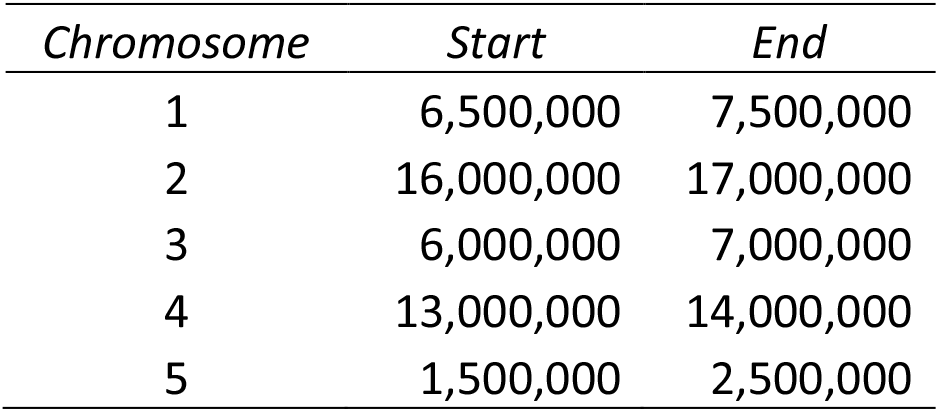
Genomic regions of *A. thaliana* in TAIR10 used for the *tsABC* and *teSMC* analyses.

| *Chromosome* | *Start* | *End* |
| --- | --- | --- |
| 1 | 6,500,000 | 7,500,000 |
| 2 | 16,000,000 | 17,000,000 |
| 3 | 6,000,000 | 7,000,000 |
| 4 | 13,000,000 | 14,000,000 |
| 5 | 1,500,000 | 2,500,000 |

