## Appendix A for "Inference of evolutionary transitions to self-fertilization using whole-genome sequences"

### Appendix A : Description of teSMC

#### 1 teSMC

To define our Hidden Markov Model (HMM) we need to define :
— Hidden States
— The signal (observed data)
— A Transition matrix (Probability of jumping from one state to another)
— An Emission matrix (Probability of observing the data given the hidden state)
— An Initial probability (Probability of hidden states at the first position of the sequence)

##### 1.1 Notations and Assumptions

We here define the different notations used and their meaning :
— $\sigma_t$ : self fertilization rate ( between 0 and 1) at time t
— $\beta_t$ : germination rate ( between 0 and 1) at time t
— $N_0$ : Population size at present time
— $r_t$ : recombination rate per nucleotide per $4N_0$ generation at time t
— $\mu$ : Mutation rate per nucleotide per $4N_0$ generation
— $\mu_b$ : ratio of mutation rate during the dormant stage over the mutation rate during the active stage per nucleotide per $4N_0$ generation
— u : time at which the recombination occurs (follows a piece-wise uniform distribution )
— L : Sequence length in bp
— $N_t$ : Population size at time t
— $\chi_t$ : Scaling factor for the population size at time t ($N_t = \chi_t N_0$)
The model's assumptions are :
— Piecewise constant population size
— Piecewise constant selfing, germination and recombination rate in time
— Constant mutation rate in time
— Constant mutation and recombination rate along the sequence
— Neutrality

##### 1.2 Hidden States

We define our hidden states at one position on the genome as the coalescent time between the two individual at that position. We note that coalescent time t (t>0). A transition from a coalescent time s to time t ($t \neq s$) at the next can only occur if a recombination happened in between the two positions.

##### 1.3 Observations

Our observations, or the signal, is a sequence of 1 and 0. This sequence is build from phasing the DNA sequences of two individual. When going along the sequence, if both nucleotide are similar, then the signal is 0 (no mutation occurred). If both are different, then a mutation occurred, and the signal is 1.

#### 1.4 Transition Matrix

A transition to state t from state s (t $\neq$ s) can only occur if there is a recombination event. Assuming Recombination event on the tree as a Poisson process we have the probability of a recombination :

$$P(rec|s) = (1 - e^{-\int_0^s \frac{2(1-\sigma_k)\beta_k}{2-\sigma_k} 2r_k dk}) \tag{1}$$

We now Assume that a recombination event occurred at time u (<s) where u follows a piece wise uniform distribution (*i.e.* uniform in each hidden state but the density between hidden state is allowed to change) between 0 and s. Then three scenarios are possible. Either the new coalescent time is smaller (t<s),bigger (t>s) or unchanged (t=s).

### 1.4.1 t<s

The resulting floating branch of the recombination event coalesces at time t < s . This mean it must not coalesce before time t (including itself). In addition we have u<t. The transition probability is therefore :

$$P(t|s,u) = \frac{2\beta_t^2}{(2-\sigma_t)\chi_t}(e^{\int_u^t -\frac{4\beta_v^2}{(2-\sigma_v)\chi_v}dv}) \tag{2}$$

### 1.4.2 t=s

The resulting floating branch of the recombination event self coalesce before time t. We therefore have the transition probability :

$$P(s|s,u) = \int_u^s \frac{2\beta_k^2}{(2-\sigma_k)\chi_k} e^{\int_u^k -\frac{4\beta_v^2}{(2-\sigma_v)\chi_v}dv} dk \tag{3}$$

### 1.4.3 t>s

The resulting floating branch of the recombination event must not coalesce (including itself) before time s. Then no coalescent event must happen before time t. We therefore have the transition probability :

$$P(t|s,u) = \frac{2\beta_t^2}{(2-\sigma_t)\chi_t} e^{\int_u^s -\frac{4\beta_v^2}{(2-\sigma_v)\chi_v}dv} e^{\int_s^t -\frac{2\beta_v^2}{(2-\sigma_v)\chi_v}dv} \tag{4}$$

##### 1.4.4 Transition probability in continuous time

In the end we have :

$$p(t|s,u) = \begin{cases} (1 - e^{-\int_0^s \frac{2(1-\sigma_k)\beta_k}{2-\sigma_k} 2r_k dk}) \frac{2\beta_t^2}{(2-\sigma_t)\chi_t}(e^{\int_u^t -\frac{4\beta_v^2}{(2-\sigma_v)\chi_v}dv}) & if \quad u<t<s \\ e^{-\int_0^s \frac{2(1-\sigma_k)\beta_k}{2-\sigma_k} 2r_k dk} + (1 - e^{-\int_0^s \frac{2(1-\sigma_k)\beta_k}{2-\sigma_k} 2r_k dk}) \int_u^s \frac{2\beta_k^2}{(2-\sigma_k)\chi_k} e^{\int_u^k -\frac{4\beta_v^2}{(2-\sigma_v)\chi_v}dv} dk & if \quad t=s \\ (1 - e^{-\int_0^s \frac{2(1-\sigma_k)\beta_k}{2-\sigma_k} 2r_k dk}) \frac{2\beta_t^2}{(2-\sigma_t)\chi_t} e^{\int_u^s -\frac{4\beta_v^2}{(2-\sigma_v)\chi_v}dv} e^{\int_s^t -\frac{2\beta_v^2}{(2-\sigma_v)\chi_v}dv} & if \quad t>s \\ 0 & if \quad otherwise \end{cases} \tag{5}$$

Once again, if all $\sigma_k = 0, \beta_k = 1$, we fall back on the probability from PSMC'.
One can find $p(t|s)$ using the total probability formula which is :

$$p(t|s) = \int_0^s p(u)p(t|s,u)du \tag{6}$$

As explained before, the state space must be finite. We therefore discretized time in n intervals. At one point the hidden state is $\alpha$ if $t \in [T_\alpha, T_{\alpha+1}]$, where $\alpha \in [0,(n-1)]$. We define $T_\alpha$ :

$$T_\alpha = -\frac{(2-\sigma_0)}{2\beta_0^2}\ln(1-\frac{\alpha}{n}) \tag{7}$$

We therefore have :

$$p(\alpha|s) = \int_{T_\alpha}^{T_{\alpha+1}} p(t|s)dt \tag{8}$$

The transition matrix need to be the probability from one state to another. Therefore we need the probability when the coalescent time at the previous position (which is here s) belongs to the state $\gamma$. To do this we simply replace s by the average coalescent time $t_\gamma$.

##### 1.4.5 Initial Probability

We use the equilibrium probability as initial probability. The equilibrium probability is the probability that the first coalescent happens in each time interval and is thus given by :

$$\begin{aligned}
q_o(\alpha) &= \int_{T_\alpha}^{T_{\alpha+1}} \frac{2\beta_\alpha^2}{(2-\sigma_\alpha)\chi_\alpha} e^{\int_0^t \frac{-2\beta_v^2}{(2-\sigma_v)\chi_v}dv} dt \\
q_o(\alpha) &= \int_{T_\alpha}^{T_{\alpha+1}} \frac{2\beta_\alpha^2}{(2-\sigma_\alpha)\chi_\alpha} e^{\int_0^{T_\alpha} \frac{-2\beta_v^2}{(2-\sigma_v)\chi_v}dv} e^{\int_{T_\alpha}^t \frac{-2\beta_v^2}{(2-\sigma_v)\chi_v}dv} dt \\
q_o(\alpha) &= e^{\int_0^{T_\alpha} \frac{-2\beta_v^2}{(2-\sigma_v)\chi_v}dv} \int_{T_\alpha}^{T_{\alpha+1}} \frac{2\beta_\alpha^2}{(2-\sigma_\alpha)\chi_\alpha} e^{\frac{-2(t-T_\alpha)\beta_\alpha^2}{(2-\sigma_\alpha)\chi_\alpha}} dt \\
q_o(\alpha) &= e^{\sum_{\eta=0}^{\alpha-1} \frac{-2\beta_\eta^2}{(2-\sigma_\eta)\chi_\eta}\Delta_\eta} (1 - e^{\frac{-2\Delta_\alpha\beta_\alpha^2}{(2-\sigma_\alpha)\chi_\alpha}})
\end{aligned} \tag{9}$$

##### 1.4.6 Calculation of $t_\gamma$

$$\begin{aligned}
t_\gamma = E[\text{Coalescent time}|\gamma] &= \frac{E[\text{Coalescent time} \cap \gamma]}{P(\gamma)} = \frac{\int_{T_\gamma}^{T_{\gamma+1}} t\Lambda_\gamma e^{-\int_0^t \Lambda_v dv} dt}{q_0(\gamma)} \\
&= \frac{\Lambda_\gamma \int_{T_\gamma}^{T_{\gamma+1}} te^{-\int_0^{T_\gamma}\Lambda_v dv} e^{-\int_{T_\gamma}^t \Lambda_v dv} dt}{q_0(\gamma)} = \frac{\Lambda_\gamma e^{-\int_0^{T_\gamma}\Lambda_v dv} \int_{T_\gamma}^{T_{\gamma+1}} te^{-\int_{T_\gamma}^t \Lambda_v dv} dt}{q_0(\gamma)} \\
&= \frac{\Lambda_\gamma \int_{T_\gamma}^{T_{\gamma+1}} te^{(T_\gamma - t)\Lambda_\gamma} dt}{(1-e^{-\Delta_\gamma\Lambda_\gamma})} = \frac{T_\gamma - T_{\gamma+1}e^{-\Delta_\gamma\Lambda_\gamma}}{(1-e^{-\Delta_\gamma\Lambda_\gamma})} + \frac{\int_{T_\gamma}^{T_{\gamma+1}} e^{(T_\gamma - t)\Lambda_\gamma} dt}{(1-e^{-\Delta_\gamma\Lambda_\gamma})} \\
&= \frac{T_\gamma - T_{\gamma+1}e^{-\Delta_\gamma\Lambda_\gamma}}{(1-e^{-\Delta_\gamma\Lambda_\gamma})} + \frac{(1-e^{-\Delta_\gamma\Lambda_\gamma})}{\Lambda_\gamma(1-e^{-\Delta_\gamma\Lambda_\gamma})} = \frac{T_\gamma - T_{\gamma+1}e^{-\Delta_\gamma\Lambda_\gamma}}{(1-e^{-\Delta_\gamma\Lambda_\gamma})} + \frac{1}{\Lambda_\gamma}
\end{aligned} \tag{10}$$

Where :

$$\begin{aligned}
\Delta_\gamma &= T_{\gamma+1} - T_\gamma \\
\Lambda_\gamma &= \frac{2\beta_\gamma^2}{(2-\sigma_\gamma)\chi_\gamma}
\end{aligned} \tag{11}$$

###### 1.4.7 Calculation of $p(\alpha|\gamma)$

$\alpha < \gamma$ We first need $p(t|t_\gamma)$ when $\alpha < \gamma$, which is obtained as described below :

$$
\begin{aligned}
p(t|t_\gamma) &= P_\gamma \int_0^t \frac{\pi_u \frac{2\beta_t^2}{(2-\sigma_t)\chi_t}\left(e^{\int_u^t -\frac{4\beta_v^2}{(2-\sigma_v)\chi_v}dv}\right)}{\Pi_\gamma} du \\
&= P_\gamma(\sum_{\eta=1}^{\alpha-1}\int_{T_\eta}^{T_{\eta+1}} \frac{\pi_u \frac{2\beta_t^2}{(2-\sigma_t)\chi_t}\left(e^{\int_u^t -\frac{4\beta_v^2}{(2-\sigma_v)\chi_v}dv}\right)}{\Pi_\gamma} du \\
&\quad + \int_{T_\alpha}^t \frac{\pi_u \frac{2\beta_t^2}{(2-\sigma_t)\chi_t}\left(e^{\int_u^t -\frac{4\beta_v^2}{(2-\sigma_v)\chi_v}dv}\right)}{\Pi_\gamma} du) \\
&= P_\gamma(\sum_{\eta=1}^{\alpha-1}\int_{T_\eta}^{T_{\eta+1}} \frac{\pi_\eta \frac{2\beta_t^2}{(2-\sigma_t)\chi_t}\left(e^{\int_u^t -\frac{4\beta_v^2}{(2-\sigma_v)\chi_v}dv}\right)}{\Pi_\gamma} du \\
&\quad + \int_{T_\alpha}^t \frac{\pi_\alpha \frac{2\beta_t^2}{(2-\sigma_t)\chi_t}\left(e^{\int_u^t -\frac{4\beta_v^2}{(2-\sigma_v)\chi_v}dv}\right)}{\Pi_\gamma} du) \\
&= P_\gamma(\sum_{\eta=1}^{\alpha-1}\int_{T_\eta}^{T_{\eta+1}} \frac{\pi_\eta \frac{2\beta_\alpha^2}{(2-\sigma_\alpha)\chi_\alpha}\left(e^{\int_u^{T_{\eta+1}} -\frac{4\beta_v^2}{(2-\sigma_v)\chi_v}dv}\right)\left(e^{\int_{T_{\eta+1}}^t -\frac{4\beta_v^2}{(2-\sigma_v)\chi_v}dv}\right)}{\Pi_\gamma} du \\
&\quad + \int_{T_\alpha}^t \frac{\pi_\alpha \frac{2\beta_\alpha^2}{(2-\sigma_\alpha)\chi_\alpha}\left(e^{\int_u^t -\frac{4\beta_v^2}{(2-\sigma_v)\chi_v}dv}\right)}{\Pi_\gamma} du) \\
&= P_\gamma(\sum_{\eta=1}^{\alpha-1}\int_{T_\eta}^{T_{\eta+1}} \frac{\pi_\eta \frac{2\beta_\alpha^2}{(2-\sigma_\alpha)\chi_\alpha}\left(e^{-(T_{\eta+1}-u)\frac{4\beta_\eta^2}{(2-\sigma_\eta)\chi_\eta}}\right)\left(e^{\int_{T_{\eta+1}}^t -\frac{4\beta_v^2}{(2-\sigma_v)\chi_v}dv}\right)}{\Pi_\gamma} du \\
&\quad + \int_{T_\alpha}^t \frac{\pi_\alpha \frac{2\beta_\alpha^2}{(2-\sigma_\alpha)\chi_\alpha}\left(e^{-(t-u)\frac{4\beta_\alpha^2}{(2-\sigma_\alpha)\chi_\alpha}}\right)}{\Pi_\gamma} du) \\
&= P_\gamma(\sum_{\eta=1}^{\alpha-1} \frac{\pi_\eta \frac{2\beta_\alpha^2}{(2-\sigma_\alpha)\chi_\alpha}\left(1 - e^{-\Delta_\eta\frac{4\beta_\eta^2}{(2-\sigma_\eta)\chi_\eta}}\right)\left(e^{\int_{T_{\eta+1}}^t -\frac{4\beta_v^2}{(2-\sigma_v)\chi_v}dv}\right)}{\frac{4\beta_\eta^2}{(2-\sigma_\eta)\chi_\eta}\Pi_\gamma} \\
&\quad + \frac{\pi_\alpha \frac{2\beta_\alpha^2}{(2-\sigma_\alpha)\chi_\alpha}\left(1 - e^{-(t-T_\alpha)\frac{4\beta_\alpha^2}{(2-\sigma_\alpha)\chi_\alpha}}\right)}{\frac{4\beta_\alpha^2}{(2-\sigma_\alpha)\chi_\alpha}\Pi_\gamma}) \\
P_\gamma &= (1 - e^{-(\sum_{\xi=1}^{\gamma-1}\frac{2(1-\sigma_\xi)\beta_\xi}{2-\sigma_\xi}2r_\xi\Delta_\xi + \frac{(t_\gamma-T_\gamma)2r_\gamma\beta_\gamma 2(1-\sigma_\gamma)}{(2-\sigma_\gamma)})})
\end{aligned} \tag{12}
$$

Where :

$$
\begin{aligned}
\pi_u &= (\frac{r_u\beta_u 2(1-\sigma_u)}{(2-\sigma_u)}) \\
\Pi_\gamma &= (\sum_{\xi=1}^{\gamma-1}\frac{\Delta_\xi r_\xi\beta_\xi 2(1-\sigma_\xi)}{(2-\sigma_\xi)} + \frac{(t_\gamma-T_\gamma)r_\gamma\beta_\gamma 2(1-\sigma_\gamma)}{(2-\sigma_\gamma)}) \\
&= (\sum_{\xi=1}^{\gamma-1}\Delta_\xi\pi_\xi + (t_\gamma-T_\gamma)\pi_\gamma)
\end{aligned} \tag{13}
$$

We can now calculate $p(\alpha|\gamma)$.

$$
\begin{aligned}
p(\alpha|\gamma) &= \int_{T_\alpha}^{T_{\alpha+1}} \frac{P_\gamma 2\beta_\alpha^2}{(2-\sigma_\alpha)\chi_\alpha}\Big(\sum_{\eta=1}^{\alpha-1} \frac{\pi_\eta(1-e^{-\Delta_\eta \frac{4\beta_\eta^2}{(2-\sigma_\eta)\chi_\eta}})(e^{\int_{T_{\eta+1}}^{t} -\frac{4\beta_v^2}{(2-\sigma_v)\chi_v}dv})}{\frac{4\beta_\eta^2}{(2-\sigma_\eta)\chi_\eta}\Pi_\gamma} \\
&\qquad + \frac{\pi_\alpha(1-e^{-(t-T_\alpha)\frac{4\beta_\alpha^2}{(2-\sigma_\alpha)\chi_\alpha}})}{\frac{4\beta_\alpha^2}{(2-\sigma_\alpha)\chi_\alpha}\Pi_\gamma}\Big)dt \\
&= \frac{P_\gamma 2\beta_\alpha^2}{(2-\sigma_\alpha)\chi_\alpha}\Big(\sum_{\eta=1}^{\alpha-1} \frac{\pi_\eta(1-e^{-\Delta_\eta \frac{4\beta_\eta^2}{(2-\sigma_\eta)\chi_\eta}})(e^{\int_{T_{\eta+1}}^{T_\alpha} -\frac{4\beta_v^2}{(2-\sigma_v)\chi_v}dv})(1-e^{-\Delta_\alpha \frac{4\beta_\alpha^2}{(2-\sigma_\alpha)\chi_\alpha}})}{\frac{4\beta_\alpha^2}{(2-\sigma_\alpha)\chi_\alpha}\frac{4\beta_\eta^2}{(2-\sigma_\eta)\chi_\eta}\Pi_\gamma} \\
&\qquad + \frac{\pi_\alpha\Big(\Delta_\alpha - \frac{(1-e^{-\Delta_\alpha \frac{4\beta_\alpha^2}{(2-\sigma_\alpha)\chi_\alpha}})}{\frac{4\beta_\alpha^2}{(2-\sigma_\alpha)\chi_\alpha}}\Big)}{\frac{4\beta_\alpha^2}{(2-\sigma_\alpha)\chi_\alpha}\Pi_\gamma}\Big)
\end{aligned}
\tag{14}
$$

$\alpha > \gamma$ We first need $p(t|t_\gamma)$ when $\alpha > \gamma$, which is obtained as described below :

$$
\begin{aligned}
p(t|t_\gamma) &= \int_0^{t_\gamma} \frac{P_\gamma \pi_u \frac{2\beta_t^2}{(2-\sigma_t)\chi_t} e^{\int_u^{t_\gamma} -\frac{4\beta_v^2}{(2-\sigma_v)\chi_v}dv} e^{\int_{t_\gamma}^{t} -\frac{2\beta_v^2}{(2-\sigma_v)\chi_v}dv}}{\Pi_\gamma} du \\
&= \frac{P_\gamma \frac{2\beta_\alpha^2}{(2-\sigma_\alpha)\chi_\alpha}}{\Pi_\gamma}\Big(\sum_{\eta=0}^{\gamma-1}\int_{T_\eta}^{T_{\eta+1}} \pi_u e^{\int_u^{t_\gamma} -\frac{4\beta_v^2}{(2-\sigma_v)\chi_v}dv} e^{\int_{t_\gamma}^{t} -\frac{2\beta_v^2}{(2-\sigma_v)\chi_v}dv} du \\
&\qquad + \int_{T_\gamma}^{t_\gamma} \pi_u e^{\int_u^{t_\gamma} -\frac{4\beta_v^2}{(2-\sigma_v)\chi_v}dv} e^{\int_{t_\gamma}^{t} -\frac{2\beta_v^2}{(2-\sigma_v)\chi_v}dv} du\Big) \\
&= \frac{P_\gamma \frac{2\beta_\alpha^2}{(2-\sigma_\alpha)\chi_\alpha}}{\Pi_\gamma}\Big(\sum_{\eta=0}^{\gamma-1}\int_{T_\eta}^{T_{\eta+1}} \pi_\eta e^{-(T_{\eta+1}-u)\frac{4\beta_\eta^2}{(2-\sigma_\eta)\chi_\eta}} e^{\int_{T_{\eta+1}}^{t_\gamma} -\frac{4\beta_v^2}{(2-\sigma_v)\chi_v}dv} e^{\int_{t_\gamma}^{t} -\frac{2\beta_v^2}{(2-\sigma_v)\chi_v}dv} du \\
&\qquad + \int_{T_\gamma}^{t_\gamma} \pi_\gamma e^{-(t_\gamma-u)\frac{4\beta_\gamma^2}{(2-\sigma_\gamma)\chi_\gamma}} e^{\int_{t_\gamma}^{t} -\frac{2\beta_v^2}{(2-\sigma_v)\chi_v}dv} du\Big) \\
&= \frac{P_\gamma \frac{2\beta_\alpha^2}{(2-\sigma_\alpha)\chi_\alpha}}{\Pi_\gamma}\Big(\sum_{\eta=0}^{\gamma-1} \pi_\eta \frac{(1-e^{-\Delta_\eta \frac{4\beta_\eta^2}{(2-\sigma_\eta)\chi_\eta}})}{\frac{4\beta_\eta^2}{(2-\sigma_\eta)\chi_\eta}} e^{\int_{T_{\eta+1}}^{t_\gamma} -\frac{4\beta_v^2}{(2-\sigma_v)\chi_v}dv} e^{\int_{t_\gamma}^{t} -\frac{2\beta_v^2}{(2-\sigma_v)\chi_v}dv} \\
&\qquad + \pi_\gamma \frac{(1-e^{-(t_\gamma-T_\gamma)\frac{4\beta_\gamma^2}{(2-\sigma_\gamma)\chi_\gamma}})}{\frac{4\beta_\gamma^2}{(2-\sigma_\gamma)\chi_\gamma}} e^{\int_{t_\gamma}^{t} -\frac{2\beta_v^2}{(2-\sigma_v)\chi_v}dv}\Big)
\end{aligned}
\tag{15}
$$

We can now calculate $p(\alpha|\gamma)$.

$$p(\alpha|\gamma) = \int_{T_\alpha}^{T_{\alpha+1}} \frac{P_\gamma \frac{2\beta_\alpha^2}{(2-\sigma_\alpha)\chi_\alpha}}{\Pi_\gamma} (\sum_{\eta=0}^{\gamma-1} \pi_\eta \frac{(1-e^{-\Delta_\eta \frac{4\beta_\eta^2}{(2-\sigma_\eta)\chi_\eta}})}{\frac{4\beta_\eta^2}{(2-\sigma_\eta)\chi_\eta}} e^{\int_{T_{\eta+1}}^{t_\gamma} -\frac{4\beta_v^2}{(2-\sigma_v)\chi_v} dv} e^{\int_{t_\gamma}^{t} -\frac{2\beta_v^2}{(2-\sigma_v)\chi_v} dv}$$
$$+\pi_\gamma \frac{(1-e^{-(t_\gamma - T_\gamma)\frac{4\beta_\gamma^2}{(2-\sigma_\gamma)\chi_\gamma}})}{\frac{4\beta_\gamma^2}{(2-\sigma_\gamma)\chi_\gamma}} e^{\int_{t_\gamma}^{t} -\frac{2\beta_v^2}{(2-\sigma_v)\chi_v} dv})dt$$
$$= \frac{P_\gamma \frac{2\beta_\alpha^2}{(2-\sigma_\alpha)\chi_\alpha}}{\Pi_\gamma} (\sum_{\eta=0}^{\gamma-1} \frac{\pi_\eta (1-e^{-\Delta_\eta \frac{4\beta_\eta^2}{(2-\sigma_\eta)\chi_\eta}})}{\frac{4\beta_\eta^2}{(2-\sigma_\eta)\chi_\eta}} e^{\int_{T_{\eta+1}}^{t_\gamma} -\frac{4\beta_v^2}{(2-\sigma_v)\chi_v} dv} + \frac{\pi_\gamma (1-e^{-(t_\gamma - T_\gamma)\frac{4\beta_\gamma^2}{(2-\sigma_\gamma)\chi_\gamma}})}{\frac{4\beta_\gamma^2}{(2-\sigma_\gamma)\chi_\gamma}})$$
$$\int_{T_\alpha}^{T_{\alpha+1}} e^{\int_{t_\gamma}^{T_\alpha} -\frac{2\beta_v^2}{(2-\sigma_v)\chi_v} dv} e^{-(t-T_\alpha)\frac{2\beta_\alpha^2}{(2-\sigma_\alpha)\chi_\alpha}})dt$$
$$= \frac{P_\gamma}{\Pi_\gamma} (\sum_{\eta=0}^{\gamma-1} \frac{\pi_\eta (1-e^{-\Delta_\eta \frac{4\beta_\eta^2}{(2-\sigma_\eta)\chi_\eta}})}{\frac{4\beta_\eta^2}{(2-\sigma_\eta)\chi_\eta}} e^{\int_{T_{\eta+1}}^{t_\gamma} -\frac{4\beta_v^2}{(2-\sigma_v)\chi_v} dv} + \frac{\pi_\gamma (1-e^{-(t_\gamma - T_\gamma)\frac{4\beta_\gamma^2}{(2-\sigma_\gamma)\chi_\gamma}})}{\frac{4\beta_\gamma^2}{(2-\sigma_\gamma)\chi_\gamma}}) e^{\int_{t_\gamma}^{T_\alpha} -\frac{2\beta_v^2}{(2-\sigma_v)\chi_v} dv} (1-e^{-\Delta_\alpha \frac{2\beta_\alpha^2}{(2-\sigma_\alpha)\chi_\alpha}})) \tag{16}$$

$\alpha = \gamma$ Because probabilities sum up to one. We have the following formula :

$$p(\gamma|\gamma) = 1 - (\sum_{\alpha=0}^{\gamma-1} p(\alpha|\gamma) + \sum_{\alpha=\gamma+1}^{n} p(\alpha|\gamma)) \tag{17}$$

#### 1.5 Emission Matrix

Because of seed banking,the coalescent tree can be very big. In this case the infinite site model hypothesis might be violated,therefore we have the following formula :

$$P(0|\gamma) = e^{-2\mu(((\beta_\gamma + ((1-\beta_\gamma)\mu_b))(t_\gamma - Tc_\gamma)) + \sum_\eta^{\gamma-1}((\beta_\eta + ((1-\beta_\eta)\mu_b))\Delta_\eta))}$$
$$P(1|\gamma) = 1 - P(0|\gamma) \tag{18}$$

Where $\mu$ is the mutation rate per nucleotide per N generation, $\mu_b$ le ratio of mutation rate during the dormant stage over the one in the active stage, $\beta_\gamma$ the germination rate in state $\gamma$ and $t\gamma$ the average coalescent time in state $\gamma$.
