## Appendix B for "Inference of evolutionary transitions to self-fertilization using whole-genome sequences"

### Appendix B – ABC method

#### Approximate Bayesian computation

Approximate Bayesian Computation (ABC) offers a statistical framework for modeling, widely used for population genetics (Beaumont, Zhang, et al., 2002). ABC addresses the two crucial modeling questions 1) which is the most supported model given the observed data, and 2) which are the most supported parameter values given the observed data from natural populations. Likelihood functions or total probabilities may be analytically complicated or computationally intractable, whereas, in the ABC framework, a posterior probability distribution is approximated using simulations. In addition, since ABC is Bayesian, it allows the implementation of prior knowledge. For this purpose, instead of using the full complexity of genetic diversity as in full-likelihood methods, the observation must be summarized into a scalar vector of summarizing statistics. The same summarizing statistics can be calculated from both simulated and natural populations. The Euclidean distance between the two simulated and observed statistics vectors determines the similarity between the simulated and true models. By defining a similarity threshold ε, we identify a set of simulations to explain the observed results. The distribution of parameters of those simulations approximates the posterior probability distribution. For this purpose, the ABC method could be optimized in a plethora of ways. The optimal performance of the ABC depends on the Bayesian sufficiency criterion of the summarizing statistic and the extensive exploration of the likelihood surface. Thus, we use PLS not only to reduce the dimensionality of the summary statistics vectors but also to reduce the influence of noise in the data. Moreover, we also apply multinomial regression (Wegmann, Leuenberger, et al. 2010) for the model choice and local linear regression for estimating the parameters. Both allow an increment of ε, causing an increase in the marginal posterior densities. This increase in the marginal density causes reduced suffering of the curse of dimensionality to improve approximation to the likelihood function.

#### Models

We propose a transition-to-selfing model (Figure 3A) and a confounding population-size model (Figure 3B) to distinguish these two effects. The transition-to-selfing model represents a diploid, random mating population of constant size, which undergoes a transition to selfing at a given time. However, the population-size model assumes a constant selfing rate.

In contrast to A, the population-size model B undergoes one single change in population size, *i. e.* we propose two possible demographic models explaining a reduced genetic diversity after a single event in the past. In both models, A and B, we define equal prior probabilities of current population sizes; however, the effective population size in the ancestral state of the scenario of A could increase to the 2-fold, only, but in B, we define the identical prior probabilities of population sizes compared to the current population size. Thus, we could identify whether a transition to selfing is inferred specifically or confounded with a simple reduction in population size.

**Simulation of genetic data**

If not indicated otherwise, from 20 individuals, we sampled 5 independent haplotypes of 1 MB length mimicking five chromosomes. For the calculation of the SFS and LD we use the 20 haplotypes. The other summary statistics are based on pairwise comparison of sequences ($TL_{true}$, $TM_{true}$, $TM_{win}$). If not indicated differently we compare all possible 20 choose 2 pairs. Thus, the total pairwise length results in 950 MB. We used mutation and recombination rates of $\mu=r=10^{-8}$. For the ABC we created 100,000 datasets for each model. We used corresponding prior parameters for both models (table S1). We tested the performance of the ABC under the assumption of neutrality. The corresponding PODs were simulated for different transition times under the shift-to-selfing model (table S1). Thus, we obtain a time series for the performance analysis. We tested the performance for following transitioning times in generations: 1,000; 2,000; 3,000; 4,000; 5,000; 6,000; 7,000; 8,000; 9,000; 10,000; 12,000; 16,000; 20,000; 30,000; 40,000; 50,000; 60,000; 70,000; 80,000; 90000; 100,000; 200,000. That translates to coalescent time units ranging from 0.05 to 10. For each condition, we created 100 independent PODs. We tested whether the assumptions of our ABC are robust if the PODs were simulated under background-selection (BGS). We used the distribution of fitness effects estimated by Hamala and Tiffin (2020) for *Arabidopsis thaliana*. We simulated sets of 5 pseudo-genomes of 1 Mb and used the annotation file of TAIR10 to determine the spatial distribution of exons. We simulated the BGS-PODs forward-in-time under Wright-Fisher assumptions. We created 100 independent burn-ins of 10*N* generations. Starting from those, we simulated a time series of transitions to explicit selfing under the same parameters as used for the neutral PODs. Each set of five simulations was aggregated and summarized into a single POD. The same set of PODS were used to test the statistical performance of *teSMC*.

**Implementation and software**

Except for BGS, we simulated the genetic data using the coalescent implemented in msprime version 0.7.4 (Kelleher, Etheridge et al. 2016). We simulated the most recent 1,000 generations under the discrete-time-Wright-Fisher to avoid biases in IBD and the following times under the SMC-prime. We used the rescaled coalescent-with-selfing (Nordborg and

Donnelly 1997) to simulate transitions to selfing for a recent selfing past. Backward-in-time, our simulations underwent a transition to outcrossing.

For the BGS-simulations, we wrote a forward-in-time simulator using *slim version 3.6* (Haller and Messer 2019). We forbid accidental selfing. We used the tree-sequence-recording option. A few lineages were not coalesced after the simulation. Thus, we recapitated the obtained tree-sequences from the *pyslim-package version 0.6*, which in turn utilizes msprime.

We implemented the whole pipeline into *Snakemake 5.13*. We have run all simulations on the high-performance cluster of the MPIPZ.

**Summarization of genetic data**

**SFS/LD**: For each sampled set of 20 haplotypes, calculated the folded SFS and the LD. The LD was calculated as $r^2$ from a subset of 10,000 randomly chosen SNPs and discretized into discrete physical distances with following breakpoints: 6,105; 11,379; 21,209; 39,531; 73,680; 137,328; 255,958; 477,066; 889,175 bp. Because recombination rates are not constant through time, we cannot relate physical distance to coalescent times. Thus, we set the discretization boundaries to fixed values.

**$TM_{win}$:** For each pairwise comparison, we counted the number of SNPs for a sliding non-overlapping window of 10,000 bp. The ABC depends on vectors of scalar summary statistics of finite size. Thus, we discretized the diversity into *m* segments, each reflecting the expected diversity under a given coalescent time. The discretization was derrived from the quantiles of the exponential distribution in order to obtain an equal information distribution per discretized diversity window via $-f * log_e \frac{1-i}{m}$. If $f = 1$, the diversity scales in coalescent times with mutation rate 1. To scale the discretization breakpoints we used $f = 8 \cdot L_{WIN} \cdot N \cdot \mu$ with $\mu$, $N, L_{WIN}$ being the per generation per bp mutation rate, the population size, and the window size, respectively. We usually choose $f$ to provide bayesian sufficiency within the time frame of expected estimates of times of transition to selfing. For the performance analysis of the ABC, we chose $N = 40{,}000$. Further*, i* and *m* were defined as $i \, \epsilon \, \{1 \ldots m\}$ *and* $m = 20$ bins. We only kept discretization breakpoints with at least a single integer value in between neighboring breakpoints. Further, we obtained the transition proportion of the $i$-th to the $(i+1)$-th window for each of the *m* discrete diversity bins. Thus, we obtained a $m^2$-transition-matrix. We flattened the matrix into a 1-dim vector of maximal length $m^2$.

**Dimensionality reduction**: The dimensionality of our summary statistics TL-distribution, $TM_{true}$, and $TM_{win}$ is up to 400 for each. To overcome the curse of dimensionality in our ABC, first, we centralized, normalized and Box Cox transformed each calculated statistic to obtain the orthogonal independent variation. Then, we applied the PLS-analysis to our data as suggested from (Wegmann, Leuenberger et al. 2010). According to the chosen combination of summary statistics, we chose an appropriate number of model selection and parameter estimation components. SFS and LD are summarizations of lower dimension. Thus, we did not necessarily apply a dimensionality reduction on them.

**Model choice**

The Bayesian framework naturally allows obtaining a posterior density for each proposed model. The Bayes factor is simply defined as the ratio of the marginal densities of two models. In ABC, we approximate those marginal densities by the posterior density. Thus, the Bayes' factor $B_{AB} = \frac{dens_A(stats_{obs})}{dens_B(stats_{obs})}$ provides an approximation to the model support given the observed data (compare Wegmann, Leuenberger et al. (2010)). We categorize the Bayes factors into negative, barley worth mentioning, substantial, strong, very strong and decisive (Jeffreys 1998).

We calculated the Bayes factor for different sets of summarizing statistics, performed a model choice using a multinomial regression analysis between proposed models A and B. The calculations were done using the R package abc (Csilléry, François et al. 2012). We accepted 1% of the total number of simulations for the model choice.

**Parameter inference**

We tested the performance of parameter inference under model A described in the Model selection section to estimate the date of a transition to selfing. We conducted the estimation using the R package abc (Csillery, Blum et al. 2010). We accepted the closest 1% of the simulations of the transitioning-model. For each corresponding set of summarizing statistics, we estimated the average posterior distribution for the 100 PODs. We show the average quantiles for the following credible intervals: 99%, 95%, 90%, 80%, 50%, 25%, 10% and the median for the whole time series. We show the performance for each parameter of the model using following sets of summary statistics: SFS/LD, $TM_{win}$, and both combined. For each we use an individual set of PLS components: No dimensionality reduction for SFS/LD, 20-PLS for $TM_{win}$, and the combined set.

**Performance / accuracy analysis**

Transitions to selfing result in a reduction of diversity. To test whether our ABC approach identifies transitions to selfing against a model of population census reduction, we conduct a model choice experiment. We approximate and compare the posterior densities of the transition to selfing model with constant population size (Figure 3C-E) to a model which undergoes a change in population size but is constant in selfing (Figure 3). We calculated the Bayes' factors using multinomial logistic regression. The results (Figure 3) depict the proportion of the correct model estimations. Depending on our summarizing statistics, our results indicate that we can precisely detect transitions to selfing for times up to 2.5 $N_e$ generations in the past if $r/\mu = 1$.

Interestingly, when using SFS/LD, we precisely detect the correct model for very recent times and outperform any other combination of parameters. However, $TM_{WIN}$ enables us to detect also older transitions to selfing. With the combination of both summarizations, we maintain a consistently high performance to model selection. The power and specificity of our model selection are more uncertain if the ratio $r/\mu$ increases.

After determining the correct model, we estimated the actual parameter of that model: When did a population undergo a transition to selfing? To test the performance of estimating the parameters, we reused the previously simulated PODs and calculated several credibility intervals for each different time point.


Beaumont, M. A. (2010). "Approximate Bayesian computation in evolution and ecology." Annual review of ecology, evolution, and systematics **41**: 379-406.

Beaumont, M. A., et al. (2002). "Approximate Bayesian computation in population genetics." Genetics **162**(4): 2025-2035.

Csillery, K., et al. (2010). "Approximate Bayesian Computation (ABC) in practice." Trends Ecol Evol **25**(7): 410-418.

Csilléry, K., et al. (2012). "abc: an R package for approximate Bayesian computation (ABC)." Methods in Ecology and Evolution **3**(3): 475-479.

Haller, B. C. and P. W. Messer (2019). "SLiM 3: Forward Genetic Simulations Beyond the Wright–Fisher Model." Molecular biology and evolution **36**(3): 632-637.

Jeffreys, H. (1998). The theory of probability, OUP Oxford.

Kelleher, J., et al. (2016). "Efficient Coalescent Simulation and Genealogical Analysis for Large Sample Sizes." PLoS computational biology **12**(5): e1004842.

Nordborg, M. and P. Donnelly (1997). "The coalescent process with selfing." Genetics **146**(3): 1185-1195.

Schiffels, S. and R. Durbin (2014). "Inferring human population size and separation history from multiple genome sequences." Nat Genet **46**(8): 919-925.

Wegmann, D., et al. (2010). "ABCtoolbox: a versatile toolkit for approximate Bayesian computations." BMC Bioinformatics **11**: 116.

Hamala T, Tiffin P. 2020. Biased Gene Conversion Constrains Adaptation in Arabidopsis thaliana. Genetics 215:831-846.
